# Beyond a Unitary Construct: Dissecting Stopping Behaviour in Two Bird Species

**DOI:** 10.1101/2024.01.17.575695

**Authors:** CA Troisi, A Vernouillet, R Allaert, S Knoch, A Martel, L Lens, F Verbruggen

**Affiliations:** Centre for Research on Cognition, Behaviour, and Ecology of Animals, Ghent University; Department of Experimental Psychology, Ghent University; Department of Pathobiology, Pharmacology and Zoological Medicine, Ghent University; Department of Biology, Ghent University

**Keywords:** inhibitory control, response inhibition, stopping of actions, race model, herring gulls, lesser black-backed gulls

## Abstract

The ability to stop behaviour is essential for adapting to changes in the environment, a principle that holds true across various species. While traditionally considered a unitary psychological construct, recent studies indicate that this ability is multifaceted. Our research evaluates this multifaceted nature using three tasks that measure stopping in different contexts in two related gull species: herring gulls (*Larus argentatus*) and lesser black-backed gulls (*L. fuscus*). These species were selected for their distinct migration and foraging strategies, offering a unique lens through which to examine behavioural adaptations. Across tasks and species, we conceptualised stopping as a race between a go and a stop runner, and predicted correlations based on the type of stop stimulus, the relative timing of the go and the stop stimuli, and the type of action that needed to be stopped. We found correlations between measures of ‘going’ across tasks, but there was less consistency in measures of ‘stopping’. Furthermore, we observed significant differences in ‘going’ and ‘stopping’ behaviours that were specific to each species, which may be linked to their migration and foraging strategies. These findings highlight the importance of considering the multifaceted nature of stopping in evolutionary and behavioural studies.

## Introduction

Inhibitory control, i.e. the suppressing or stopping of actions and thoughts, is widely considered a critical component of flexible and adaptive behaviour (Diamond, 2013). It allows individuals to act with restraint, quickly alter their behaviour, and solve new problems (V. J. Brown & Tait, 2014; Diamond, 2013; Griffin et al., 2016; Mettke-Hofmann, 2014), which can have significant fitness consequences (e.g. Minter et al., 2017; Moffitt et al., 2011). Take for example a bird wanting to forage on some seeds it spotted on the ground. Just before leaving the bushes, it detects a cat jumping from a nearby wall, but fails to stop the action of going towards the seeds and gets predated. In this example, not having been able to stop the action ultimately led to a major fitness loss for the bird (but an increased one for the cat!).

In the current study, we focused on the action component of inhibitory control, which we will refer to as ‘stopping (of actions)’ throughout the rest of this paper (for reviews on the different types of inhibition, see Bari & Robbins, 2013; Nigg, 2000). Drawing upon research from different domains, we predicted correlations based on the type of stop stimulus (e.g., the cat in the previous example), its relative timing (e.g., when the cat was detected by the bird), and the type of action that needs to be stopped (e.g., flying towards the seeds). We then tested these predictions in an experiment focusing on two closely related gull species that performed three different stopping tasks. A better understanding of how stopping is constructed will allow us to make more accurate predictions about mechanisms, causes (e.g. developmental differences) and consequences (e.g. fitness) of individual variation in stopping behaviour (Verbruggen et al., 2014; Völter et al., 2018).

Many researchers seem to assume (explicitly or implicitly) that stopping of actions happens similarly in different contexts, which is also reflected in the fact that various tasks purporting to measure stopping of actions are often used interchangeably within- and across studies. Across these tasks, stopping can be described as an independent race between a ‘go runner’, which is triggered by a ‘go stimulus’ (e.g. a piece of food), and a ‘stop runner’, which is triggered by a ‘stop stimulus’ (e.g. a predator suddenly appearing) (Logan & Cowan, 1984). Whether stopping is successful or not will depend on the relative finishing time of the runners: if the go runner finishes first, the action will be executed (i.e. stopping is unsuccessful); by contrast, when the stop runner finishes first, the action will be stopped (i.e. stopping is successful). The race model has been successfully used to describe stopping across modalities and species (e.g. hand movements in humans, eye movements in monkeys, lever presses or nose pokes in rodents, whole body movements (walking) in sheep, or pecking in pigeons) (Eagle & Robbins, 2003; Hanes & Schall, 1995; Knolle et al., 2017; Lea et al., 2019; Logan & Cowan, 1984; for reviews, see: Schall & Godlove, 2012; Verbruggen & Logan, 2009), making it popular across domains. The broad applicability of the race model seems, at first sight, consistent with the idea that stopping is a unitary concept. However, while the race model provides a good description of behavioural outcomes (in terms of relative finishing times), it does not provide a description of the go and stop runners themselves. While it is generally accepted that there can be many differences in the go runner (e.g. type of action, such as pecking a seed vs. flying towards a patch), the same level of diversity may exist for the stop runner across tasks or situations.

Detailed analyses have indeed revealed differences in stop runners in terms of, e.g. the stimulus that triggers the stop runner in the race, the moment this stop runner can be triggered (relative to the go runner), and the action that must be stopped (Bari & Robbins, 2013; Beran, 2015; Bray et al., 2014; Brucks et al., 2017; Hervault et al., 2021; Littman & Takács, 2017; Munakata et al., 2011; Swick et al., 2011; Van Belle et al., 2014; Verbruggen et al., 2014; Verbruggen & Logan, 2008a; Völter et al., 2018). First, stopping may be influenced by the stop stimulus. Often, stopping is triggered by external stimuli. This can be a sudden or salient change in the environment (e.g. a red traffic light for humans, or a predator for a bird). However, the stop stimulus could also be the overall context. For instance, humans typically don’t check their phones during meetings. Similarly, a low-ranking animal might wait with eating until higher-ranking animals have left the food patch. Furthermore, in some situations there may be no external stop stimuli at all and stopping is triggered by an internal stimulus, such as a change in motivational state or a conflict between different (action) options. While research on humans (and a few animal species) indicates that stopping in response to external vs. internal stimuli engages only partly overlapping neural networks (Ridderinkhof et al., 2014; Van Belle et al., 2014), how stopping across tasks might differ as a function of the characteristics of the stimulus that triggers the stopping has received relatively little attention in the animal cognition domain (but see Dewulf et al., in prep; Zucca et al., 2005). Furthermore, sudden or salient stimuli (e.g. a loud sound) might also trigger an initial global pause (followed by cancelation of the initial action) that is absent for contextual or internal stop stimuli (Diesburg & Wessel, 2021). Second, stopping may be influenced by the relative timing of the go and stop stimuli (Sebastian et al., 2013; Swick et al., 2011). For example, when a bird spots a nut, it may plan to fly towards it, but may then stop at the last minute when it suddenly spots a predator. In this example, there is a delay between the presentation of the go stimulus (i.e. food) and the stop stimulus (i.e. the predator). By contrast, if a bird has learned that it can eat brown nuts but not similarly shaped brown pebbles, it will eventually peck at one class of brown shapes (nuts) and not peck when it encounters another class of shapes (pebbles). In this example, there is no delay between the go and stop stimuli. It has been argued that stopping in such a situation (nut vs pebble) will take place at decision or selection stages (targeting specific actions), whereas stopping in the former situation (food and predator) will involve a different ‘global’ stopping mechanism (suppressing all motor output) because the go runner is already initiated before the stop stimulus appears (Littman & Takács, 2017; Munakata et al., 2011; Rubia et al., 2001; Swick et al., 2011). Third, stopping may be influenced by the very nature of the to-be-stopped action. As mentioned above, the race model applies to different behaviours (Verbruggen & Logan, 2009). But while this is the case for the stopping of single ‘discrete’ actions (such as going towards a feeder), a different picture seems to emerge for the stopping of ‘repetitive’ actions (such as perseverative pecking at the feeder that is covered), with recent work showing that both might be associated with different neural signatures (Hervault et al., 2021; Wadsley et al., 2022).

## Unravelling Variability Across Tasks

The above review of the stopping literature suggests that stopping actions consists of different subcomponents. This could explain why many animal cognition studies found no (or only low) correlations between different tasks that purportedly measure stopping (Anderson et al., 2017; Boogert et al., 2011; Bray et al., 2014; Brucks et al., 2017; Shaw et al., 2015; Troisi et al., 2021; van Horik, Langley, Whiteside, Laker, & Madden, 2018; van Horik, Langley, Whiteside, Laker, Beardsworth, et al., 2018; Vernouillet et al., 2018; Völter et al., 2022; but see Ashton et al., 2018; Davidson et al., 2022; Montalbano et al., 2020; Sollis et al., 2022).

The first aim of the present study was to study variability in stopping behaviour across tasks. We therefore used three different tasks to study stopping, namely a detour barrier task, a thwarting task, and a stop-change task. In the following sections, we describe each task (as used in the present study) and which subcomponents it may measure (Table 1). Based on this, we make predictions about correlations between the behavioural measures across tasks.

**Table 1:**
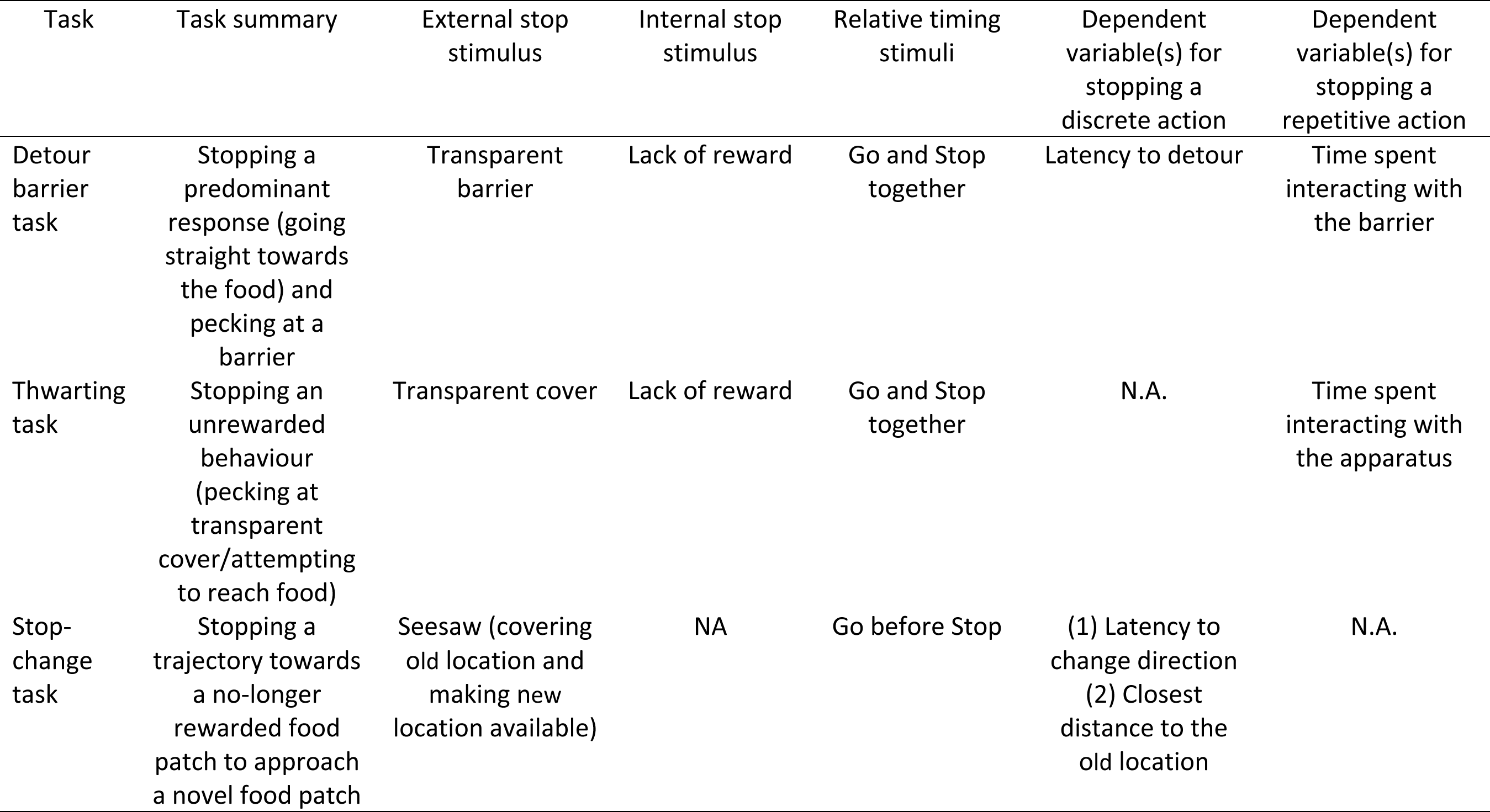
Overview of the three tasks used in our study, including a task summary, possible external and internal stop stimuli, relative timing of go and stop stimuli, and type of action that has to be stopped (with the corresponding dependent variables). For all tasks the go stimulus was the presence of food, and the dependent variable measured for going was the latency to interact with the task. See Method section for details.

### Detour barrier task

Individuals were faced with a transparent barrier, behind which there was visible food. They had to suppress the response to run straight towards the food and instead detour around the barrier. The interpretation of the go component of the task is straightforward, as the go runner is triggered by the visible food presented behind the barrier. However, interpreting the stop component is less straightforward. The stop runner in the race could be triggered by various external, and possibly, internal stimuli. First, the barrier itself may have acted as an external stop stimulus (Kabadayi et al., 2018) as the individuals had previous experience with transparent barriers in their home enclosures. Second, the overall context and test arena may have acted as an external contextual stop stimulus (Kabadayi et al., 2017), as the test arena strongly resembled the feeding stations in the home enclosures, where the food was hidden behind non-transparent barriers. Additionally, each individual had direct prior experience with the test arena itself, again with non-transparent barriers as part of their training. Third, the retrieval of previous ‘detour’ memories (during training with an opaque barrier) may have acted as an internal stop stimulus (Wallis et al., 2001; for a similar idea in other stopping tasks see, e.g. Verbruggen & Logan, 2008b) as the animals were trained to detour to obtain food. In these three scenarios, the relative timing of go and stop stimuli is expected to be the same (i.e. there is no delay between the presentation of the go and stop stimuli). Finally, the task involves (at least initially) stopping a discrete (single) action, namely running towards the food.

When individuals failed to stop the initial response to run straight towards the food and instead started pecking the barrier, the detour barrier task also measured a second component of stopping. That is, to obtain the reward, the individual first had to stop the ongoing but unrewarded action (i.e. pecking at the barrier) (e.g. van Horik, Langley, Whiteside, Laker, Beardsworth, et al., 2018). As noted above, stopping repetitive actions, such as perseverative pecking, may be distinct from stopping the initial response to run towards the food (Hervault et al., 2021; Wadsley et al., 2022). This stopping of the repetitive action could be triggered by an external stimulus (i.e. the barrier or context; see above) or an internal stimulus, related to the non-delivery of the reward. In both scenarios, the go stimulus (the food behind the barrier) and stop stimulus are present simultaneously.

### Thwarting task

A familiar food bowl was placed in the centre of the test arena and covered with a transparent lid, making the food visible but inaccessible (except for a single piece of fish placed on top of the cover). The go runner is presumably triggered by the presence of the food bowl and the accessible piece of fish. In terms of stopping: the transparent cover may have acted as an external stimulus if individuals generalized their experiences with transparent barriers (from their home enclosures and the detour barrier task) to the transparent cover. Stopping pecking or interacting with the food bowl may also be triggered by an internal stop stimulus, similar to stopping pecking at the barrier in the detour barrier task. In both cases, the relative timing of go and stop stimuli is the same, and individuals have to stop a repetitive action (i.e., pecking or trying to access the food underneath the cover).

### Stop-change task

Food was initially visible at one location in the test arena. When the individual approached the food, the location of the food unexpectedly changed using a seesaw (see the Methods section). Here, the go runner is again triggered by the presentation of the food (clearly visible at a specific location in the test arena). The stop runner is triggered by an external stop stimulus, namely the seesaw and the accompanying change in food location. Unlike in the other two tasks, during the stop-change task, the external stop stimulus appears after the go stimulus. Individuals have to stop a discrete action (going towards the previously visible food location) in this task.

### Across-task correlations

In each task, the go runner is triggered by the presentation of food. Reactions to this go stimulus can be driven by motivation, general processing speed, or aspects of personality such as activity and exploration (Carere & Locurto, 2011; Dougherty & Guillette, 2018; Miyake et al., 2000; Sih & Del Giudice, 2012; Troisi et al., 2019). Therefore, we expect correlations between go behaviour in each task. In both the thwarting and stop-change tasks, we have a relatively straightforward measures of ‘going’. In the thwarting task, this corresponds to the time between entering the test arena and interacting with the food bowl for the first time, and in the stop-change task, this corresponds to the time between entering the test arena and triggering the seesaw (when the individual was halfway towards the visible food location). In both cases, short latencies indicate a fast go runner and are therefore expected to correlate with each other. In the detour barrier task, we measure the time needed to interact with the task for the first time (which is either the first peck at the barrier or detouring around the barrier, whichever comes first). However, detouring around the barrier is less pure as a measure of going, as it could be influenced by stopping as well. This could weaken the correlation with the measures of going in the other tasks.

Regarding ‘stopping’, we expected correlations between some stopping measures, but not necessarily others, depending on the overlap between the task components (See Table 1 and Figure 1). In terms of stop stimuli and their relative timing, the detour barrier and thwarting tasks were more similar to each other than to the stop-change task: both tasks have similar external and internal stop stimuli (i.e. transparent objects and non-delivery of reward, respectively), and there is presumably no delay between the presentation of go and stop stimuli. By contrast, a different stop stimulus is used in the stop-change task (a seesaw), which appears well after the go stimulus (and after the ‘go runner’ has already been initiated). In terms of the nature of the actions that had to be stopped: the detour barrier and stop-change tasks both involved stopping a discrete single action (i.e. running towards the food). In the stop-change task, we can directly measure this stop-change latency (the latency between the time the bird triggers the seesaw, and the time it changes direction), as well as the measure of distance of the bird from the unrewarded location. In the detour barrier task, we use the latency to successfully detour as a measure of stopping the response to go straight (though this measure is again less pure than the one obtained in the stop-change task, as going and stopping cannot be disentangled). Furthermore, if stopping the initial response in the detour barrier task failed, it also measured the stopping of a repetitive action (perseverative interacting with the barrier), akin to the time the bird spent interacting with the (covered) food bow during the thwarting task.

**Figure 1:**
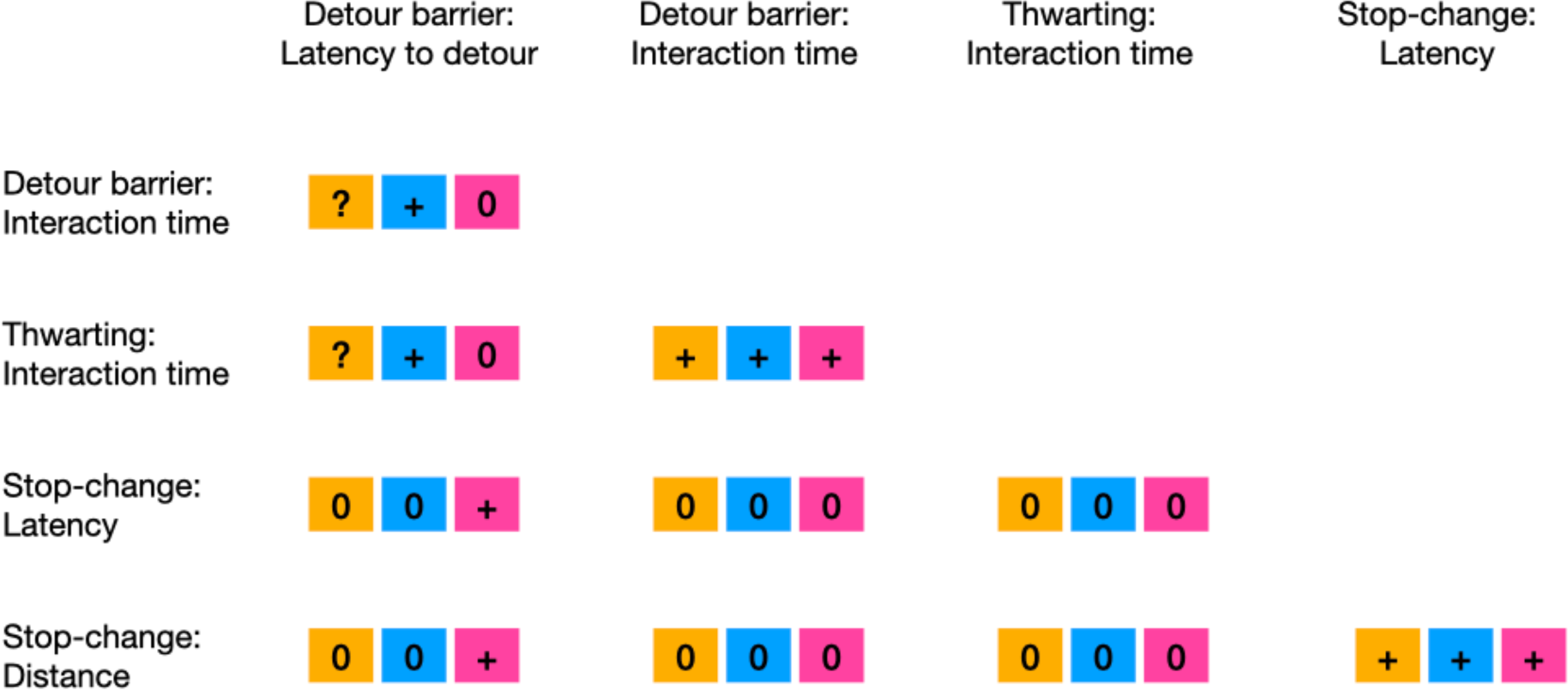
Predicted correlations between different measures of stopping (top row and first column; see main text for description) based on the overlap between task components (i.e. type of stop stimulus, orange; relative timing of go and stop stimuli, blue; type of action that has to be stopped, purple). **+** indicates we predicted a correlation; **(0)** indicates we predicted no correlation. Note that for the detour barrier task, we could not always make a priori predictions (indicated by **?**) as the initial stop stimulus in this task is unclear (see the description of the detour barrier task for further details); but once they pecked at the barrier, the stop stimuli in the detour barrier task would be similar to the stop stimuli in the thwarting task (i.e. a transparent object or the failure to obtain a reward). Measures of ‘going’ were not included in this figure, but correlations among these across tasks were predicted. We did not make any predictions about correlations between ‘going’ and ‘stopping’ measures.

## Unravelling Variability Across Species

A second aim of this study is to study variability in stopping for two ecologically and phylogenetically related species, the herring gull (*Larus argentatus*) and the lesser black-backed gull (*L. fuscus*) (Kim & Monaghan, 2006). Both species demonstrate high flexibility (both between and within individual) in their use of the environment (Belant, 1997; Rock & Vaughan, 2013; Spelt et al., 2019, 2021; Tyson et al., 2015), making them suitable model species for studying the stopping of actions. But despite the many similarities, herring gulls and lesser black-backed gulls also exhibit some key species differences in, e.g. migration and foraging strategies. We explored if such differences are associated with variability in one or more stopping subcomponents. Building further on the race model, we also explored measures of going.

Unlike lesser black-backed gulls, herring gulls are not long-distance migrants. Therefore, herring gulls will have to adjust their foraging strategies (e.g. different food sources, foraging techniques) to changes in resource availability over time, stopping to use previously rewarding foraging patches or techniques. We might, therefore, expect herring gulls to be more efficient at stopping than lesser black-backed gulls in response to changes in the *immediate* environment (Mettke-Hofmann, 2010). The two species also differ in their food resources and strategies to access them. For example, compared with lesser black-backed gulls, herring gulls dig more for food when feeding on refuse (Verbeek, 1977), and they tend to feed more in intertidal zones where they must also dig for food (Garthe et al., 1999; Kim & Monaghan, 2006; Sotillo et al., 2014). As such, we could speculate that herring gulls will perseverate more in situations when the food is not immediately accessible (i.e. they will take longer at stopping a repetitive action that does not immediately lead to a reward). Thus, even though we could not make strong predictions about the direction of the effects, we had good reasons to assume that the herring gulls and lesser black-backed gulls differ in at least some stopping components.

## The Present study

The current study had two main aims. First, it aimed to explore the variation in stopping behaviour by using three unique tasks, each designed to probe different aspects of stopping as a function of the type of stop stimulus, the relative timing of the go and stop stimuli, and the type of action being stopped. Secondly, the study compared the stopping behaviour of two closely related bird species, the herring gull and the lesser black-backed gull. The aim of this comparison was to understand whether their different migratory and foraging behaviour was related to their ability to stop. Through this dual approach, the study aimed to provide insights into both the task-dependent nature of stopping behaviour and its variation between species.

## Methods

Detailed information, following the MeRIT system (Nakagawa et al., 2023), about all methods and procedures is provided in the Supplementary Materials. We used Large Language Models for proofreading.

### Subjects

#### Egg collection and incubation

From May 2021 to June 2021, eggs were collected by the Agency for Nature and Forests (ANB) and the Wildlife Rescue Centre Ostend (WRC) and brought to the WRC on the day of collection. Upon arrival, the eggs were weighed, measured, and photographed before incubating them. This was done until we reached our target sample size of 120 (Table S1).

#### Chick rearing

After hatching, chicks were kept indoors. They were moved to outdoor enclosures (10 m^2^) when they were approximately 5 days old. Each enclosure held 15 chicks of similar age (except for the last two enclosures where individuals had up to 13 days of age difference). Originally, we aimed to rear chicks under predictable and unpredictable conditions (see Supplementary Materials). However, technical problems during incubation delayed the study, and prevented us from implementing the early-life manipulation as planned. Therefore, we included predictability treatment as a control variable rather than an experimental variable in our between-species analyses.

After testing, when individuals were between 25 and 39 days old, they were housed in a large flight cage (approximately 180 m^2^) for approximately four to six weeks (depending on the finishing time of the tests) and were subsequently released in the wild.

#### Species ID and sex

Species ID and sex were confirmed through DNA sampling, from down feathers collected on the day of hatching. If DNA sampling was not possible for an individual, we identified their species using morphological characteristics when they were ringed and predicted their sex with a support vector machine classifier using morphological data (see Supplementary Materials for a validation of this method).

### Behavioural tests

#### Group habituation and training in the home enclosure

There were two feeding stations per enclosure, in which food was placed behind opaque barriers. This provided chicks with detour experience (Figure 2). In addition, three transparent barriers (50 x 100 cm width x height) were placed within the non-feeding area of the enclosure. These barriers provided chicks with experience with transparency. Both the transparent and opaque barriers had coloured tape on the sides to delimit the area of the barrier.

**Figure 2:**
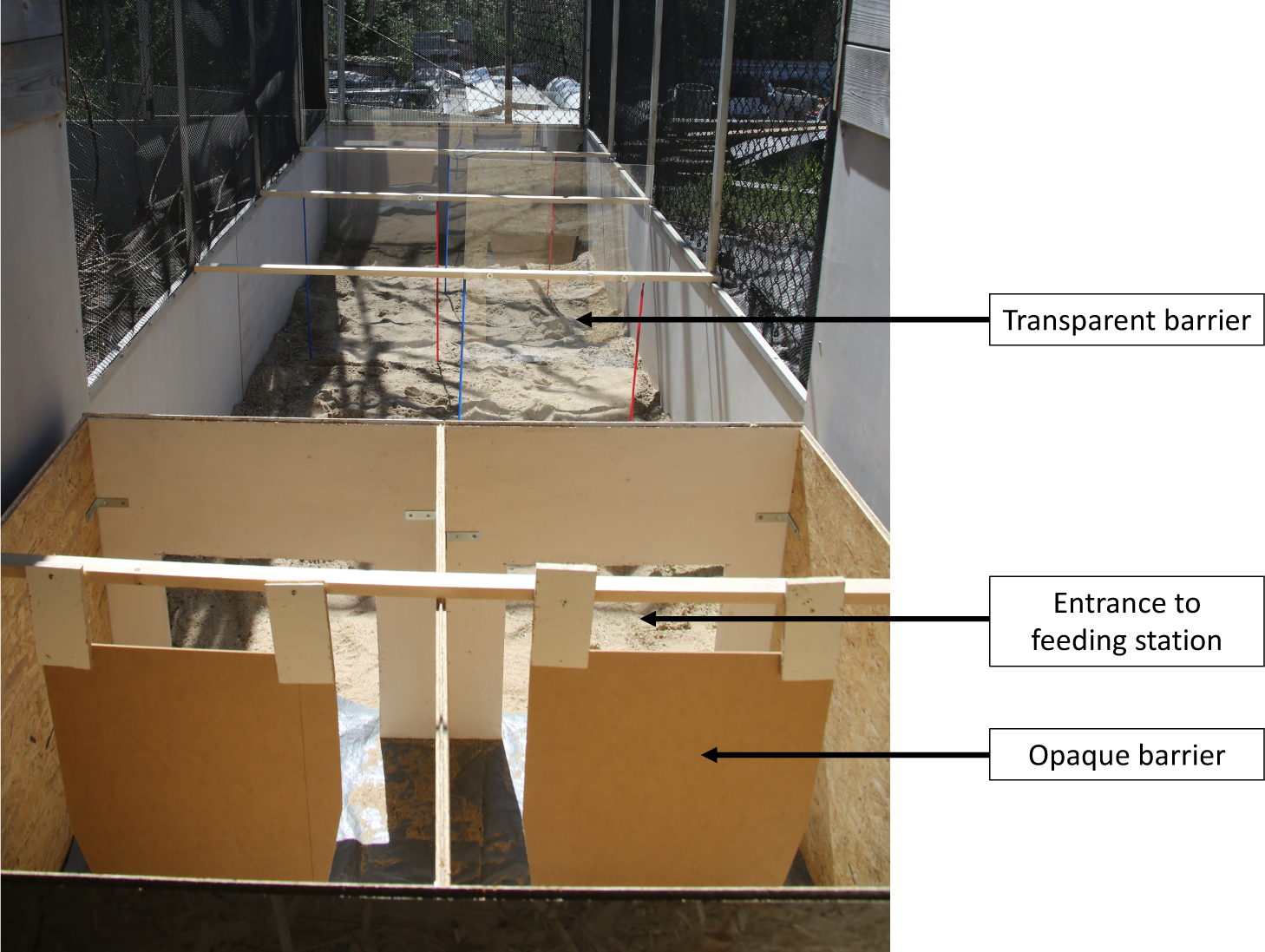
Enclosure with feeding station with opaque barriers in the foreground, and three sets of transparent barriers in the background. Note: not pictured here, but opaque barriers also had coloured tape around their edges, in a similar way to the transparent barriers.

#### General testing protocol

Behavioural tests started 7 days after the group was complete. See Figure 3 for an overview. Mean age on the first day of testing was 16.7 days (range 13-21 days; due to human error, the exact hatching date was unknown for 19 individuals). Two enclosures (of similarly aged chicks) were tested each day (one enclosure from each predictability treatment; see above). For each testing day, birds were food deprived at 18:00 the previous day and were tested in the morning (8:00-11:00). Due to human error, the birds of one enclosure received food prior to testing on the 7^th^ July 2021 (stop-change task), while on 16^th^ July 2021 (also stop-change task), birds of another enclosure were not food deprived in the evening (but were not given food in the morning). Note that the stop-change task took place a few days after the other tests for practical reasons.

**Figure 3:**
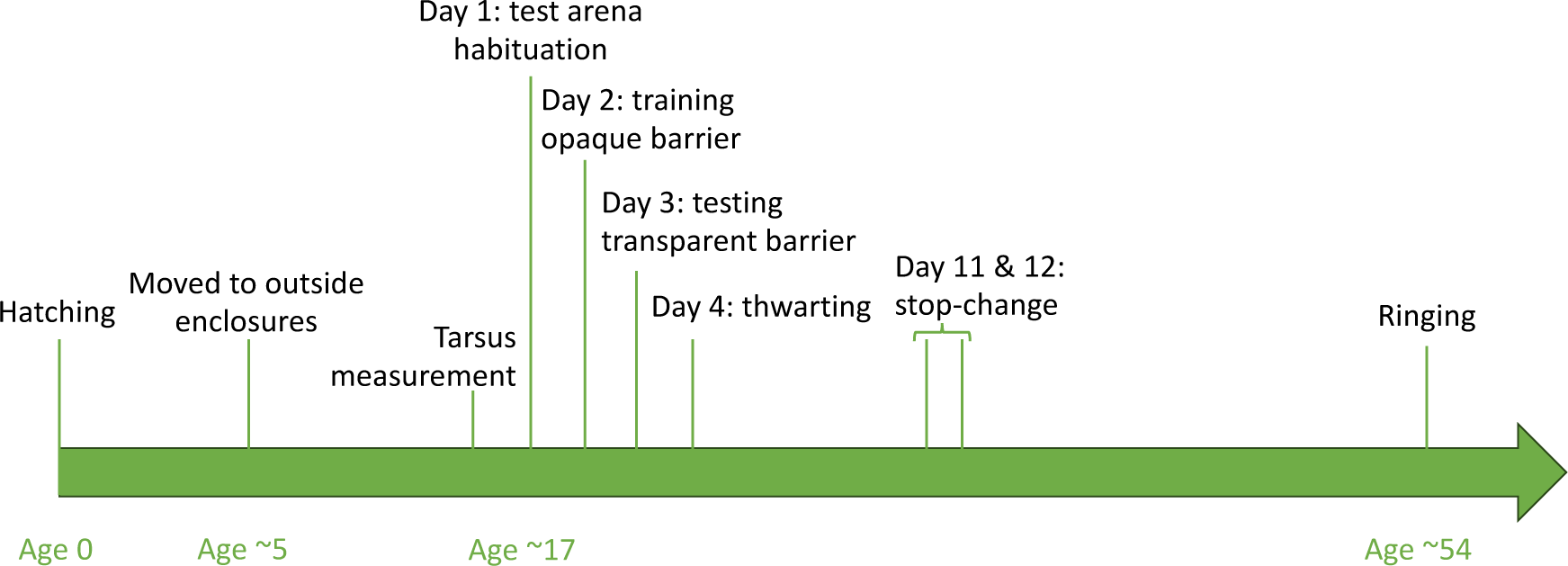
Timeline of the experiment, including mean age of the birds in days

Behavioural tests were conducted in two test boxes, equipped with cameras. Individuals were transported from their enclosure to these boxes in a cat carrier. They were then placed in a start box connected with a sliding door to the main test arena (Figure 4). Unless stated otherwise, individuals were left in the start box for 30 s, before the door between the start box and the test arena was opened. At this point, the trial started. If birds did not exit the start box within 60 s, they were gently pushed forward with the back of the start box sliding forward. Trials ended either when individuals reached the food (for the detour barrier and stop-change tasks) or when the time limit of the trial was reached (all tasks). Individuals could not see the experimenter during testing. At the end of the trial, individuals were put back in their cat carrier, and placed in a dark room. The order of testing was semi-random: experimenters picked the first bird that they came across within the enclosure. Once all individuals finished testing, they were all placed back in their enclosure and fed.

**Figure 4:**
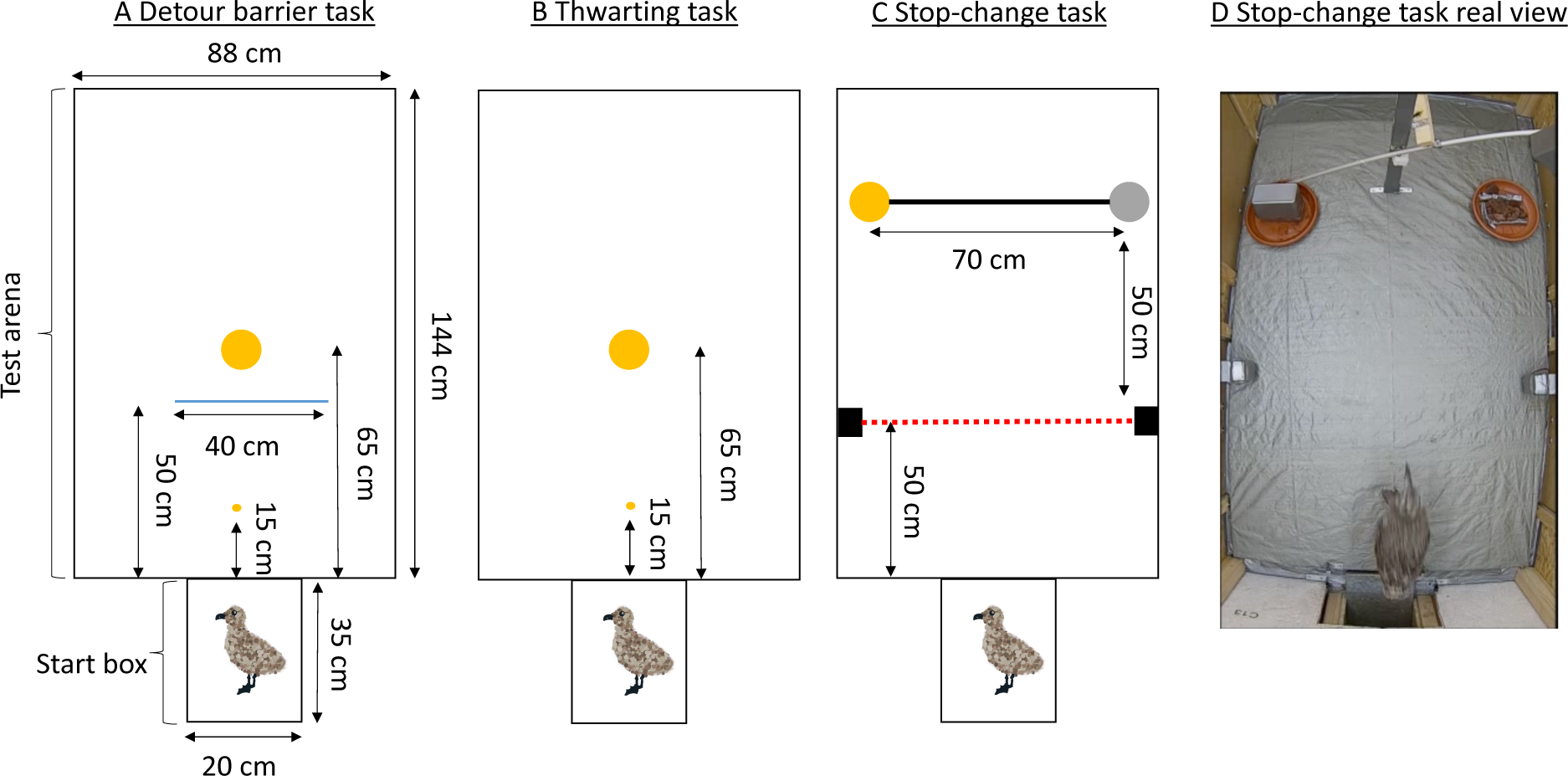
Schematics of the three tasks used, including dimensions (height start box: 26 cm, test box: 132 cm): (A) detour barrier task, (B) thwarting task, (C) stop-change task. The large yellow circles represent food, and the grey circle in the stop-change task represents food that is made non-available during the task. The small yellow circles (in the detour barrier and thwarting task) represent the start food (see main text). The blue line in the detour barrier task represents the (opaque and transparent) barrier. The red dotted line in the stop-change task represents the infra-red beam. Gull drawings by AV. (D) is a real view of the stop-change task at the start of the trial, with the food on the left being covered, and the food on the right being initially accessible.

#### Individual habituation in test arena

On Day 1, chicks were individually habituated to the test arena. A food bowl (diameter: 17 cm) containing fish was placed 50 cm from the entrance, and a small piece of fish was placed in front of the start box entrance (henceforth “start food”; 15 cm from the entrance). This was included to measure motivation, but we noticed during testing that many individuals ignored this start food and immediately ran towards the main food bowl. Therefore, consumption of this start food was not used as a measure of motivation in our analyses. Once the door between the start box and test arena was opened, birds were left for 300 s in the test arena.

#### Detour barrier task

On Day 2 (individual detour training), an opaque barrier (40 * 40 cm length * height, made of cardboard, with coloured tape on each side delimiting the barrier) was placed 50 cm from the start box entrance, in front of a bowl of food (65 cm from the start box entrance; see Figure 4A). Start food was again placed at the entrance of the test arena. Once the door between the start box and test arena was opened, birds were given a max of 180 s to detour the barrier and eat the food placed behind the barrier. On Day 3 (detour test), a similar set up to the individual training was used, except the barrier was made of transparent plastic. The trial stopped once the individual had eaten, or after 180 s after the start of the trial.

#### Thwarting task

On Day 4, individuals took part in the thwarting task. A food bowl (17 cm diameter), covered by a transparent plastic sheet (rendering the food visible but inaccessible), was placed 65 cm from the entrance of the start box. A piece of food was placed 15 cm from the entrance, as well as on top of the covered food bowl (Figure 4B). After opening the door between the start box and the test arena, individuals had 180 s to interact with the inaccessible food. After 180 s, the transparent cover over the food bowl was removed, making the food accessible. Individuals had 60 s to interact with the now-accessible food.

#### Stop-change task

On Days 11 and 12, the stop-change task took place. The apparatus consisted of a seesaw with two cups, and an infrared beam triggering the seesaw. The infrared beam was placed 50 cm from the entrance of the start box, and the food bowls (with cups) were placed 50 cm from this beam (Figure 4C). At the start of the trial, the cup on the right was approximately 50 cm above the food bowl, while the cup on the left was covering the food. Food bowls were 70 cm apart (Figure 4C, 4D). The seesaw was held in place by an electromagnet. Upon breaking the infrared beam, the magnet would switch off, allowing the seesaw to tilt, covering the food on the right (henceforth “old location”), and uncovering the food on the left (henceforth “new location”). Unlike in the other tasks, the start box had a transparent door, allowing individuals to see the location of the food before the start of the trial (always to the right, Figure 4D). The birds were placed in the start box for 15 s, after which the door was opened. If individuals did not exit after 30 s, they were gently pushed forward into the test arena. The trial stopped once the individual had eaten some food, or 120 s after they had entered the test arena.

### Dependent variables and inclusion criteria

Behaviour during all tasks was video recorded, and videos subsequently coded using BORIS (Friard & Gamba, 2016). The video coder was naive to the hypotheses, treatment, and to the species and sex of the individual. 20% of videos were coded by a second coder (naïve to the species, treatment, and sex of the individual, but not naïve to the hypotheses) to calculate inter-rater reliability.

#### Measures of going

For all tasks, we used the time between leaving the start box (i.e. when the chick’s feet were outside the box) and the first interaction with the task as the measure of going. For the detour barrier task, the first interaction was either the time they came in physical contact with the barrier (usually through pecking), or for those that directly detoured the barrier without interacting with it, the latency to detour (see below). For the thwarting task, the first interaction was either pecking at the free piece of fish on the bowl or pecking at the cover itself. For the stop-change task, we considered the crossing of the infrared beam as the first interaction with the task.

#### Measures of stopping

In the detour barrier task, we had two measures of stopping. First, we measured the latency to detour, which refers to the time between the moment of leaving the start box (i.e. when the chick’s feet were outside the box) and the moment the chick’s feet crossed the (virtual) line of the barrier. Second, we measured the time spent physically interacting with the barrier. For the thwarting task, we also measured the time spent physically interacting with the (covered) bowl. Finally, for the stop-change task, we used the stop-change latency as our primary measure of stopping. This was defined as the time between crossing the infrared beam and the moment the chick changed direction (i.e. orientating the body towards the new location instead of the old location). Consistent with previous work (Meier et al., 2017), we also considered the spatial characteristics of the ‘change point’ as a secondary measure. The smallest distance between the chick and the old location was calculated using the Tracker software (D. Brown et al., 2012).

#### Control variables

Previous work suggests that stopping is influenced by the general motivational state or activity level of the individual (e.g. van Horik, Langley, Whiteside, Laker, Beardsworth, et al., 2018). Therefore, we also included an independent measure of motivation or activity in our between-species comparison. For this, we measured the latency to exit the start box, which is defined as the time between opening the door of the start box and the moment the chick’s feet are outside the start box. The measures of going and stopping could also be influenced by the size of the chick. Therefore, we also measured tarsus length measured the day before the habituation session. We used an average of two measures of the left tarsus, and two measures of the right tarsus, taken at the same time.

#### Inclusion criteria

In the analyses reported below, we only included chicks that ‘participated’ in the tests (Table S2). In the detour barrier task, this was defined as interacting with the barrier or eating food behind the barrier (N=99); in the thwarting task, this was defined as interacting with the covered food bowl or eating food once the cover was removed (N=105); and in the stop-change task, this was defined as crossing the infrared beam (N=106). We had to omit 7 individuals from the stop-change task because of technical issues, resulting in a sample size of 99. For the ‘Variability Across Tasks’ analyses in which we included all tasks, the sample size is n=87 (i.e., individuals that participated in all three tasks). For the analysis of the latency to interact with the stop-change task in the ‘Variability Across Species’ analysis, the model had a convergence issue, which was fixed by removing a clear outlier (n=98).

Excluding individuals could have introduced STRANGE biases (Webster & Rutz, 2020) as our final sample might not have been representative of the average individual. However, we believe this to be unlikely as we found that species ID, treatment (manipulated over enclosures), sex, and tarsus length did not significantly predict task participation (Tables S3-5).

#### Ethical statement

We performed the experiment in accordance with the Association for the Study of Animal Behaviour ethical guidelines under permission of the ethical committee of animal experimentation (VIB Site Ghent, Universiteit Gent; EC2021-017). Eggs were collected under permit ANB/BL-FF/V21-00154.

#### Data Availability

Data and R Code are available on OSF: https://osf.io/jbe4q/?view_only=cd763b15f4b649fb80e520fea326f0a3.

## Statistical Analysis

All analyses were conducted in R version 4.2.0 (R Core Team, 2022). Inter-coder reliability was assessed using the interclass correlation coefficient from the *icc* function from the *irr* package (version 0.84.1, Gamer et al., 2019); consensus between the two coders was high (Table S6). The packages *ggplot2* (version 3.4.3, Wickham, 2016), *jtools* (version 2.2.2, Long, 2022) and *cowplot* (version 1.1.1, Wilke, 2020) were used for plotting graphs. We used a ‘co-pilot’ system (Reimer et al., 2019) to check the data processing and data analysis code.

### Variability Across Tasks

As a first step, to assess the relationship between pairs of measures, as hypothesised and outlined in Figure 1, we performed correlations between the behavioural measures using the *cor* function in the *stats* package (R Core Team, 2022). This approach allowed us to assess the strength and direction of linear relationships between each pair of variables, providing an initial understanding of the components across tasks. As we also predicted that some measures would not correlate with each other, we calculated Bayes Factors using the *correlationBF* function in the package *BayesFactor* (version 0.9.12-4.6, Morey & Rouder, 2023). We used the default prior width as used in R for BayesFactors. A Bayes Factor > 1 is in favor of the alternative hypothesis, whereas a Bayes Factor < 1 is in favor of the null hypothesis. A Bayes Factor around 1 yields inconclusive evidence. The size of the Bayes Factor determines whether the evidence is anecdotal (1/3 − 1; 1-3), moderate (1/3 − 1/10; 3-10), strong (1/10 − 1/30; 10-30), very strong (1/30 − 1/100; 30-100) or extreme (< 1/100; > 100) (Schönbrodt & Wagenmakers, 2018).

In a second step, to further determine whether the data suggested ‘unity’ or ‘diversity’ across multiple measures and tasks, we used Principal Component Analysis (PCA). This analysis was based on the correlation matrix using all stopping measures together and was performed using the *prcomp* function in the *stats* package (R Core Team, 2022). The results of the PCA were interpreted to see whether a small number of components could explain a significant proportion of the variance in the data, suggesting ‘unity’, or whether the variance was more evenly distributed across several components, suggesting ‘diversity’. It is important to note that we didn’t predict species-specific changes in the relationship between the different go and stop components. Therefore, we collapsed the data from all individuals for these two sets of analyses. This decision was based on the understanding that the first main aim of our study was to capture general trends across species, rather than to explore species-specific differences.

For the ‘Variability Across Tasks’ analyses, missing values (e.g. when an individual interacted with the apparatus but did not detour, we could not calculate the detour latency) were replaced with the time the individual spent in the testing arena. This corresponds to the maximum duration of the trial minus the time spent in start box (Table S2). Note that for the detour barrier task, there were 16 individuals (out of 99) for which we used the same measure for the first interaction with the task and the latency to detour (because they did not peck at the barrier). We therefore reran all analyses after excluding those birds. The results are reported in the supplementary material (Table S7-9; Figure S1) and the main results remain similar, except that we find anecdotal rather than moderate evidence of a correlation between the latency to interact with the detour task and the latency to interact with the stop change task, and we find very strong rather than moderate evidence of a correlation between the two measures of stopping in the stop-change task (Table S7).

### Variability Across Species

Given that we did not find one main component of stopping (see PCA results below), we analysed each behavioural measure separately to explore if the two species varied in going and stopping. We corrected for multiple comparisons within each task (for the detour barrier and stop-change task, we had three measures, so alpha=0.017; for the thwarting task, we had two measures, so alpha=.025).

We included the following fixed effects: species (herring gulls vs lesser black-backed gulls), treatment (predictable vs unpredictable), sex (female vs male), and (scaled) latency to exit the start box as a general measure of motivational state and activity level. We included sex because previous studies found sex differences in tasks measuring stopping (Lucon-Xiccato, 2022). We included enclosure as a random effect. For measures that involved running towards the food (i.e. latency to interact with the apparatus, detour latency, stop-change latency, and distance), we included (scaled) tarsus length to control for morphological differences.

We rounded our measures to obtain count data (time or distance) and started with the Poisson family model. If model assumptions were violated, we then tried negative binomial distribution. If model diagnostics showed evidence of zero-inflation or heteroskedasticity, we also included a zero-inflated model and a dispersion model. Model families for each model are stated in the Supplementary Material. For linear mixed models we used the *glmer* function from the *lme4* package (version 1.1-34, Bates et al., 2015) and for negative binomial models we used the *glmmTMB* function from the *glmmTMB* package (version 1.1.7, Brooks et al., 2017). Model assumptions were checked using the *DHARMa* package (version 0.4.6, Hartig, 2020).

## Results

### Variability Across Tasks

#### Correlations

For going, we found moderate support for a positive correlation between the latency to interact for the first time with the detour barrier task and the equivalent measure in the stop-change task (*r* = 0.261, *BF* = 4.04; Figure S2). However, we found only moderate support for a correlation between the latency to interact for the first time with the detour barrier task and that of the thwarting task (*r* = 0.206, *BF* = 1.38), and moderate support for the *absence* of correlation between this latency in the thwarting and the stop-change tasks (*r* = 0.073, *BF* = 0.304; Figure S2).

The correlations for measures of stopping are shown in Table 2. We found that there was moderate support for a correlation between latency to detour and latency to change direction in the stop-change task (Table 2, Figure S2). We also found extreme and moderate support for within-task correlations in the detour barrier and stop-change tasks, respectively (Table 2, Figure S2). Finally, we found moderate support for the *absence* of a correlation between time spent interacting with the barrier in the detour barrier and (a) the time spent interacting in the thwarting tasks and (b) the latency to change in the stop-change task. There was also moderate support for the absence of a correlation between latency to detour and distance to the old location in the stop-change task.

**Table 2:** Correlation matrix showing the correlation coefficient and Bayes Factor (in brackets) between the different behavioural measures of stopping (N=87). In bold regular are results with moderate to extreme support for the alternative hypothesis, and in bold italic are results with moderate support for the null hypothesis.

|  | Detour barrier:<br>Latency to<br>detour | Detour barrier:<br>time spent<br>interacting with<br>the barrier | Thwarting: time<br>spent interacting<br>with the<br>apparatus | Stop-change:<br>latency to<br>change |
| --- | --- | --- | --- | --- |
| Detour barrier:<br>time spent<br>interacting with<br>the barrier | **0.571 (1573001)** |  |  |  |
| Thwarting: time<br>spent interacting<br>with the<br>apparatus | -0.171 (0.800) | **0.039 (0.261)** |  |  |
| Stop-change:<br>latency to<br>change | **0.283 (6.78)** | **-0.068 (0.295)** | -0.154 (0.636) |  |
| Stop-change:<br>minimum<br>distance to old<br>location | **0.071 (0.302)** | -0.100 (0.368) | -0.185 (0.988) | **0.277 (5.85)** |

#### PCA

The results are shown in Table 3, and Figure 7. Axis 1 had positive loadings for the latency to interact with the stop-change task (going), the latency to detour in the detour barrier task (stopping), and the latency to change in the stop-change task (stopping). Axis 2 had negative loadings for the detour latency (stopping) and for the time spent interacting with the detour barrier (stopping), but a positive loading for the distance to the old location during the stop-change task (stopping). Finally, Axis 3 had a negative loading for the latency of the first interaction in the thwarting task (going) and a positive loading for the time spent interacting with the apparatus in the same task (stopping).

**Figure 7:**
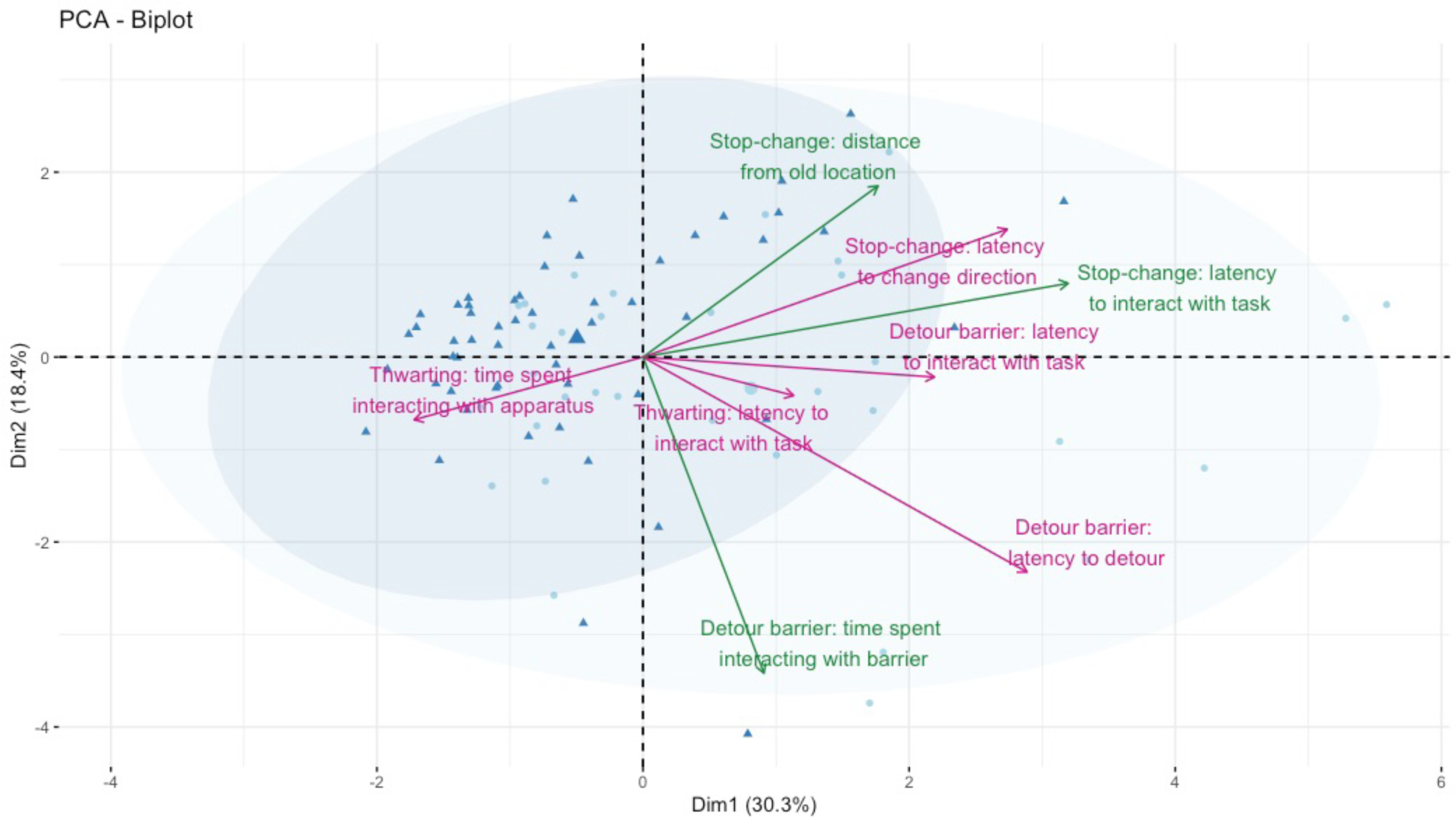
Biplot of the first two principal component analysis axis, which together explain 48.7% of the variation in stopping behaviour across the detour barrier, thwarting and stop-change task. Herring gull individuals are labelled in light blue, and lesser black-backed gulls are labelled in dark blue. In green are variables related to the go component, while in pink are variables related to the stopping component.

**Table 3:** Structure of the principal component analysis for behavioural measures related to going and stopping behaviour in the detour barrier, thwarting and stop-change tasks (N=87). We only report the three axis whose variance is bigger than predicted if each variable contributed equally to the variance (the full PCA is reported in Table S11). In bold we report the loading for variables whose loading is bigger than expected if all variables contributed equally to the specific axis. We ordered the variable according to the subcomponent it is related to: we first present the three latencies to interact with the task (going), then the five measures related to stopping.

|  | Axis 1 | Axis 2 | Axis 3 |
| --- | --- | --- | --- |
| Std deviation | 1.56 | 1.21 | 1.09 |
| Proportion of variance | 0.303 | 0.184 | 0.149 |
| Cumulative proportion | 0.303 | 0.487 | 0.636 |
| Detour barrier: latency to interact with the task (Go) | 0.351 | -0.044 | -0.270 |
| Thwarting: latency to interact with the task (Go) | 0.181 | -0.085 | **-0.699** |
| Stop-change: latency to interact with the task (Go) | **0.511** | 0.164 | 0.249 |
| Detour barrier: latency to detour (Stop) | **0.462** | **-0.477** | 0.101 |
| Detour barrier: time spent interacting with the barrier (Stop) | 0.146 | **-0.702** | 0.161 |
| Thwarting: time spent interacting with the apparatus (Stop) | -0.275 | -0.139 | **0.487** |
| Stop-change: latency to change (Stop) | **0.438** | 0.283 | 0.311 |
| Stop-change: minimum distance to old location (Stop) | 0.283 | **0.380** | 0.075 |

#### Variability Across Species

Given the lack of clear overarching stopping components, we analysed each behavioural measure separately to examine variability across species. Lesser black-backed gulls were significantly faster to first interact in the detour barrier (Table 4A, Figure 8A), thwarting (Table 4B, Figure 8B) and stop-change tasks (Table 4C, Figure 8C) than herring gulls. They were also significantly faster to detour in the detour barrier task (Table 4D, Figure 8D) and change direction in the stop-change task (Table 4E, Figure 8E). For the latency to interact with the detour, the latency to detour in the detour barrier task and the latency to change direction in the stop-change task, we found that lesser black-backed gulls were also less variable than herring gulls (Tables S12-14, Figure 8). See Supplementary Materials for the other analyses in which we did not find species differences (Tables S15-20).

**Figure 8:**
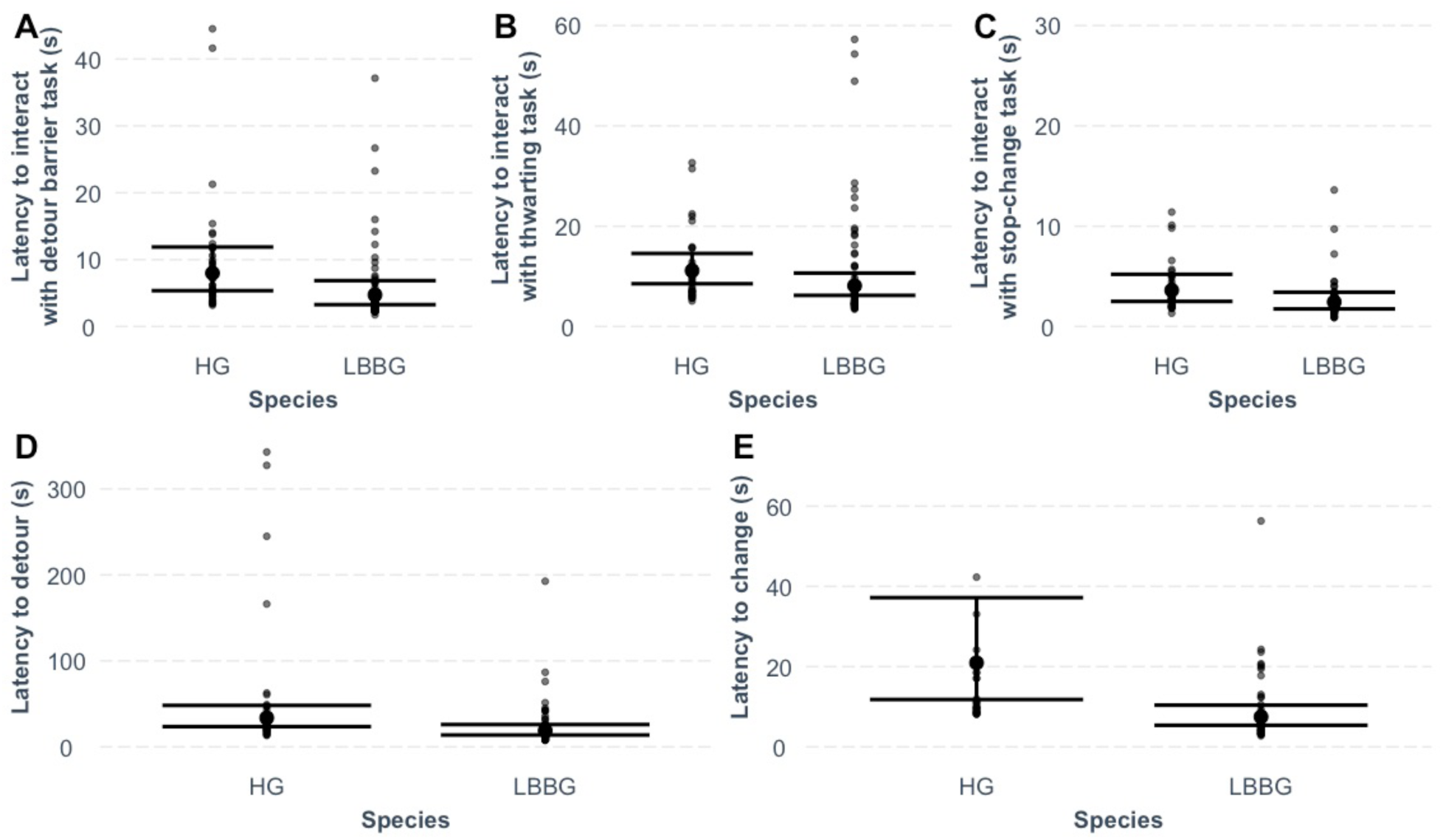
Partial residual plot showing species differences in (A) the latency to interact with the detour barrier task (s) (n=99), (B) the latency to interact with the thwarting task (s) (n=103) (C) the latency to interact with the stop-change task (s) (n=98), (D) the latency to detour (s) in the detour barrier task (n=93), and (E) the latency change (s) in the stop-change task (n=82). Error bars are 95% confidence intervals.

**Table 4:**
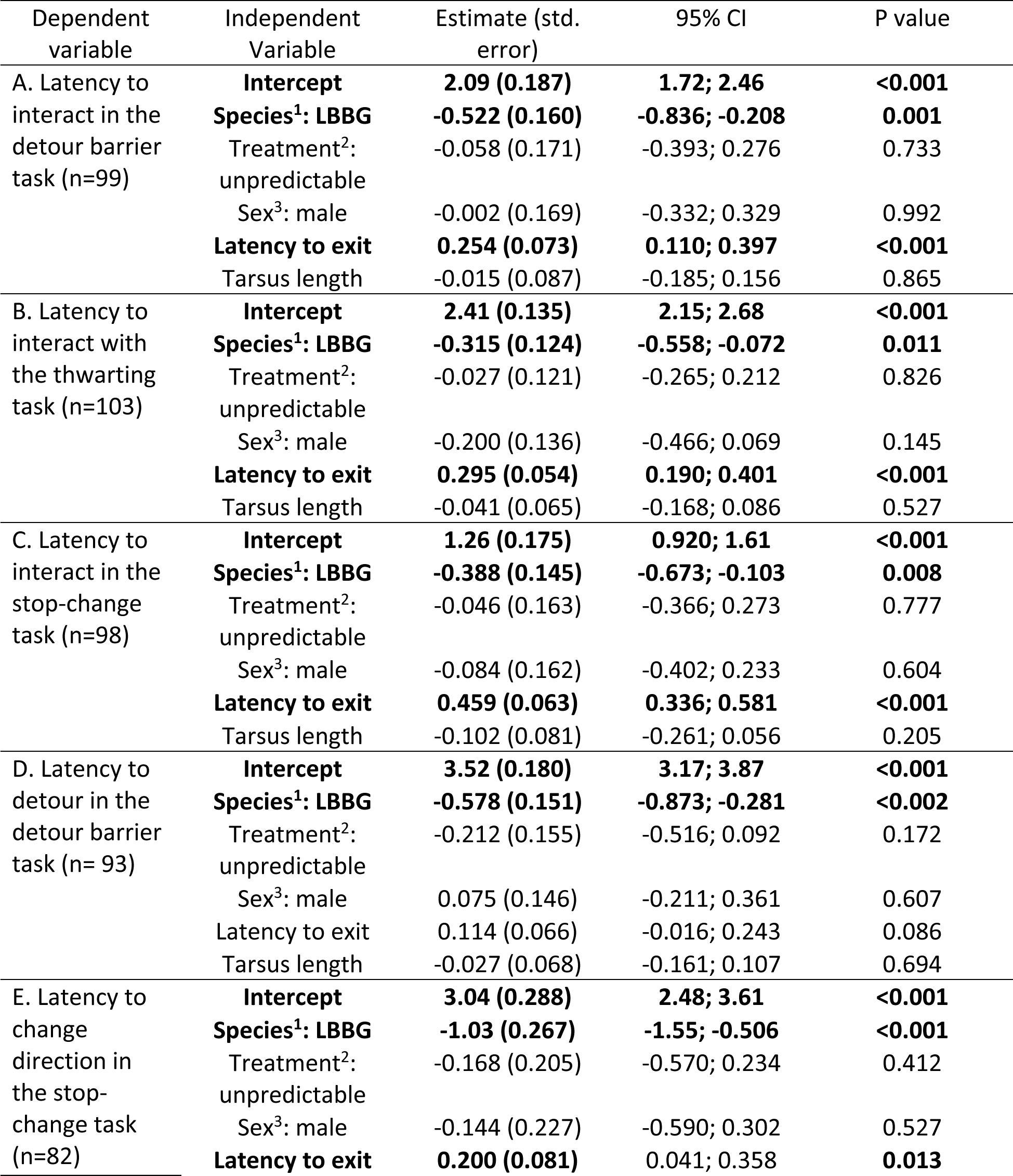
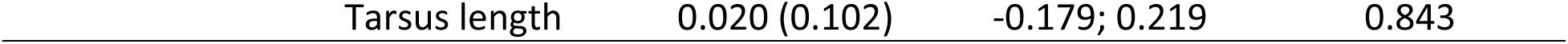
Models showing the effect of species, treatment, sex, latency to exit, and average tarsus length on latency of first interaction in the detour barrier task (A), thwarting task (C) and stop-change task (C); latency to detour in the detour barrier task (D); and latency to change direction in the stop-change task (E). Enclosure was always included as a random effect. For all measures, we used a negative binomial model accounting for overdispersion. The dispersion models included species, treatment and sex, as main effects. ^1^ baseline = herring gull; ^2^ baseline = predictable treatment; ^3^ baseline = female

| Dependent variable | Independent Variable | Estimate (std. error) | 95% CI | P value |
| --- | --- | --- | --- | --- |
| A. Latency to interact in the detour barrier task (n=99) | **Intercept** | **2.09 (0.187)** | **1.72; 2.46** | **<0.001** |
|  | **Species<sup>1</sup>: LBBG** | **-0.522 (0.160)** | **-0.836; -0.208** | **0.001** |
|  | Treatment<sup>2</sup>: unpredictable | -0.058 (0.171) | -0.393; 0.276 | 0.733 |
|  | Sex<sup>3</sup>: male | -0.002 (0.169) | -0.332; 0.329 | 0.992 |
|  | **Latency to exit** | **0.254 (0.073)** | **0.110; 0.397** | **<0.001** |
|  | Tarsus length | -0.015 (0.087) | -0.185; 0.156 | 0.865 |
| B. Latency to interact with the thwarting task (n=103) | **Intercept** | **2.41 (0.135)** | **2.15; 2.68** | **<0.001** |
|  | **Species<sup>1</sup>: LBBG** | **-0.315 (0.124)** | **-0.558; -0.072** | **0.011** |
|  | Treatment<sup>2</sup>: unpredictable | -0.027 (0.121) | -0.265; 0.212 | 0.826 |
|  | Sex<sup>3</sup>: male | -0.200 (0.136) | -0.466; 0.069 | 0.145 |
|  | **Latency to exit** | **0.295 (0.054)** | **0.190; 0.401** | **<0.001** |
|  | Tarsus length | -0.041 (0.065) | -0.168; 0.086 | 0.527 |
| C. Latency to interact in the stop-change task (n=98) | **Intercept** | **1.26 (0.175)** | **0.920; 1.61** | **<0.001** |
|  | **Species<sup>1</sup>: LBBG** | **-0.388 (0.145)** | **-0.673; -0.103** | **0.008** |
|  | Treatment<sup>2</sup>: unpredictable | -0.046 (0.163) | -0.366; 0.273 | 0.777 |
|  | Sex<sup>3</sup>: male | -0.084 (0.162) | -0.402; 0.233 | 0.604 |
|  | **Latency to exit** | **0.459 (0.063)** | **0.336; 0.581** | **<0.001** |
|  | Tarsus length | -0.102 (0.081) | -0.261; 0.056 | 0.205 |
| D. Latency to detour in the detour barrier task (n= 93) | **Intercept** | **3.52 (0.180)** | **3.17; 3.87** | **<0.001** |
|  | **Species<sup>1</sup>: LBBG** | **-0.578 (0.151)** | **-0.873; -0.281** | **<0.002** |
|  | Treatment<sup>2</sup>: unpredictable | -0.212 (0.155) | -0.516; 0.092 | 0.172 |
|  | Sex<sup>3</sup>: male | 0.075 (0.146) | -0.211; 0.361 | 0.607 |
|  | Latency to exit | 0.114 (0.066) | -0.016; 0.243 | 0.086 |
|  | Tarsus length | -0.027 (0.068) | -0.161; 0.107 | 0.694 |
| E. Latency to change direction in the stop-change task (n=82) | **Intercept** | **3.04 (0.288)** | **2.48; 3.61** | **<0.001** |
|  | **Species<sup>1</sup>: LBBG** | **-1.03 (0.267)** | **-1.55; -0.506** | **<0.001** |
|  | Treatment<sup>2</sup>: unpredictable | -0.168 (0.205) | -0.570; 0.234 | 0.412 |
|  | Sex<sup>3</sup>: male | -0.144 (0.227) | -0.590; 0.302 | 0.527 |
|  | **Latency to exit** | **0.200 (0.081)** | **0.041; 0.358** | **0.013** |

The above analyses (Table 4) revealed a general effect of motivational state and activity level. Specifically, we found that individuals that were faster at exiting the start box were also significantly faster at interacting for the first time in the detour barrier, thwarting and stop-change tasks (Table 4.A-C), and significantly faster at stopping in the stop-change task (Table 3.E). In the thwarting task, they also spent significantly more time pecking at the cover (Table S19). Finally, in the stop-change task, they were significantly closer to the old location (Table S20). Thus, general motivational state or activity level influenced some of the measures of going and stopping. By contrast, tarsus had no significant effect on any of the measures, and sex significantly influenced only one measure: males were less variable in their latency to first interact with the thwarting apparatus than females (Table S15).

To further examine whether any species differences observed in our going and stopping behaviour might be underlined by species difference in motivation or activity, we also investigated whether there were any species differences in the latency to exit the start box (Tables S21-24). Latency to exit the start box was repeatable across tasks (r= 0.326, 95% CI = 0.241, 0.417, p<0.001, n=120; only birds that participated in the three tasks: r= 0.283, 95% CI = 0.181, 0.389, p<0.001, n=87; see supplementary material) and for individuals that participated in the tasks, lesser black-backed gulls were faster at exiting the start box compared to herring gulls in the stop-change task (Table S25).

## Discussion

The first aim of the study was to investigate, in two phylogenetically and ecologically related gull species, whether different subcomponents of stopping behaviour across three different tasks were related to each other. Our results show that, in line with previous work in the human cognitive psychology, neuroscience literature, and animal cognition field, stopping an action is not a unitary construct (Bari & Robbins, 2013; Beran, 2015; Diamond, 2013). The second aim of the study was to investigate whether these gull species differ in going or stopping components. Species differences, especially in migratory behaviour, have received some interest in relation to personality (Mettke-Hofmann, 2010, 2014, 2017; Mettke-Hofmann & Gwinner, 2004; Nilsson et al., 2010). The results of our study suggest that differences in migration and foraging strategies may also influence aspects of cognition.

## Unravelling Variability Across Tasks

We conceptualised stopping of actions as a race between a go runner and a stop runner. Given that all tasks involved a similar go stimulus (a food reward) and action (approaching the food), we expected that the measures of going would correlate across tasks. By contrast, we expected differences across stopping components, as the tasks differed (at least party) in terms of stop stimuli, the relative timing of the go and stop stimuli, and the type of action that had to be stopped.

### Going

Unlike what we predicted, our measures of going were not correlated across all tasks. While we found moderate support for a positive correlation between our measure of going in the detour barrier task and going in the stop-change task, we did not find such support for a correlation between the measure of going in the thwarting task and the corresponding measures in the detour barrier and stop-change tasks.

The lack of consistent correlations between the thwarting task and the other two tasks could be due to the lower variability in behaviour in the thwarting task (Figure S1), which may have reduced the statistical power. This reduction in variability could be explained by the fact that in the thwarting task, the bowl was presented in the centre of the test box. This meant that, unlike in the detour barrier and stop-change tasks, there was only one location where individuals could interact with the task. Previous work has shown that the spatial properties of a task could influence individual performance (Troisi et al., 2021). In addition, we placed a small piece of fish on top of the bowl, which could have made the go stimulus even more salient, further reducing variability.

The correlation between the detour barrier and stop-change task measures could be due to several factors, such as individual differences in general motivation for food, anxiety, cognitive processing speed (e.g. food detection or decision making), or even walking speed. The present study does not allow us to distinguish between these different possibilities. Nevertheless, our results suggest that there is some ‘unity’ across tasks as far as going is concerned.

### Stopping

The findings from the stop-change and detour barrier tasks indicate that the type of action that needs to be stopped matters (i.e. stopping discrete single actions, such as running towards a food location, vs. stopping repetitive actions, such as perseverative pecking at a cover). Specifically, we observed in these tasks moderate support for a positive correlation between two measurements of stopping discrete actions, namely the detour latency and stop-change latency, even though the stop stimuli and relative timing of the stimuli differed between tasks (see also Figure 1). Both latency measures were also associated with the same principal component (Axis 1). Based on these findings, we could also expect a correlation between the distance to the old location in the stop-change task and the latency in the detour barrier task. We did not observe such a correlation; in fact, we even found moderate support for the null hypothesis. As mentioned above, the absence of the predicted correlation could be explained by the impurity of the ‘distance’ measure, as it includes both components of going and stopping. The ‘distance’ measure, and the latency to detour do however load together on PCA Axis 2. Finally, we also found moderate to extreme support for within-task correlations between the measures of stopping in the detour-barrier and stop-change tasks. It should be noted, however, that the within-task measures were not completely independent. That is, the latency to detour is likely to increase as the individual pecks longer at the barrier. Similarly, an individual that stops and changes quickly, is also more likely to remain further away from the old location than an individual that stops and changes slowly (assuming, that they were initially running equally fast).

The analysis of the latency measures suggests a degree of overlap in the stopping of discrete actions in detour barrier and stop-change tasks. However, the latency to interact with the stop-change task, which we used as a measure of the ‘going’, also strongly loaded on the same PCA axis (Table 3, Figure 7). This suggests that the correlation between the measures of stopping discrete actions might not solely be attributable to a commonality in stopping. Thus, similar to the across-task correlations in going (see previous section), the across-task correlations in stopping might be (at least partly) due to individual variation in, for example, general cognitive processing speed, motivation, activity level, anxiety, or exploration tendencies (Brucks et al., 2017; Carere & Locurto, 2011; Dougherty & Guillette, 2018; Miyake et al., 2000; Munakata et al., 2011; Rozas et al., 2008; Sih & Del Giudice, 2012).

If the nature of the to-be-stopped action matters, one could also expect a correlation between two specific measures of stopping repetitive actions: the duration of interacting with the barrier in the detour barrier task and with the covered food bowl in the thwarting task. van Horik et al. (2018) found such a correlation in pheasant chicks (*Phasianus colchicus*). We could not replicate this correlation (and even found moderate support for the null hypothesis). As summarised in Table 1, the detour barrier and thwarting tasks shared various features. However, there were still some differences that may explain the lack of correlation. First, the individuals in our study were familiar with transparent objects similar to those in the detour barrier task from their home enclosure. This prior experience could have influenced their ability to recognise and navigate around the barrier more effectively (Stow et al., 2018; van Horik, Langley, Whiteside, Laker, Beardsworth, et al., 2018). Second, the thwarting task featured a free piece of fish on the cover of the food bowl, a feature absent in the detour barrier task. This could have acted as a partial reinforcement, potentially influencing the pecking behaviour. Third, the variability in stopping in the thwarting task was smaller, which, like the measure of going behaviour, might have obscured potential correlations with the pecking in the detour barrier task.

Another possibility for the absence of some (expected) correlations is that some stopping measures were less reliable than others. Although we measured multiple (sub)components of stopping, we had only one trial per task. Determining the reliability and repeatability of cognitive measures requires multiple trials. However, task performance is strongly influenced by learning, which could in turn influence the measure of repeatability. Indeed, in detour tasks, individuals tend to improve and become faster over trials (reviewed in Kabadayi et al., 2018). To reduce the effect of learning, one could, for example, introduce different types of barriers during the detour barrier task (Davidson et al., 2022; Dewulf et al., in prep; McCallum & Shaw, 2023; Sollis et al., 2022), different bowls and covers in the thwarting task, or different locations in the stop-change task. Nevertheless, the issue remains that consistency observed across trials may be due to factors that are unrelated to stopping.

In sum, our research suggests that the nature of the action to be stopped plays a more critical role in eliciting consistent behaviour across tasks than the exact stop stimulus and the relative timing of the go and stop stimuli, at least in the detour barrier and stop-change tasks.

### The race model and implications for the study of stopping

Whether individuals can stop or not ultimately depends on the race between the go and stop runners (Logan & Cowan, 1984). It is therefore crucial to consider both aspects together. Despite the popularity of the detour barrier task, disentangling measures of going and stopping in this task is challenging for various reasons. First, it is unclear which stimulus triggers the stop runner. It is often assumed to be the barrier, but it could also be another external or internal stimulus (see Introduction), making it difficult to determine *if* and *when* stopping is initiated. Second, since stopping can theoretically be initiated at any point in time, obtaining a pure (latency) measure of going is also not straightforward. Finally, it follows from the race model that the outcome (whether they detour or not) cannot be used as a pure measure of stopping either, because it depends on both going and stopping.

In the stop-change task, we can independently measure the initial go response (i.e. the time needed to run from the start box towards the point of the infrared beam, triggering the seesaw) and the subsequent stop-change response (i.e. the time needed to stop and change the response after the seesaw has dropped). This allows for at least a partial dissociation between the going and stopping components, which is a significant advantage of this task. In fact, the stop-change task as used here was originally designed with the race model in mind, building on existing and well-established tasks in the (human) inhibitory control and stop-signal literature (Verbruggen & Logan, 2009; Verbruggen & McLaren, 2017).

However, it is important to recognise that although the race model assumes that the go and stop runners run ‘independently’ most of the time, it does not exclude the possibility that both runners are similarly influenced by the same underlying factors (De Jong et al., 1990; Verbruggen & Logan, 2009). Thus, general (non-inhibitory) factors, such as the ones discussed above (e.g. motivation, personality, processing speed), could still contribute to overall task performance (faster going and stopping) and explain correlations between going and stopping (both within- and between-tasks). Crucially, as the race model is not a process model, it does not necessarily explain the origin of correlations between stopping across tasks. To do so, we need both precise behavioural measures (something the race model can be used for) and careful manipulations of tasks and contexts (e.g. by manipulating the stop stimulus within the same task).

Across domains, the race model has been used to explain the stopping of a discrete action. Whether it also applies to the stopping of repetitive actions remains an open question. In any case, the lack of between-task correlations suggest that it is harder to measure than measuring the stopping of a discrete action. This could be due to the lack of an obvious external stop stimulus. Performance in the thwarting task might also be (even) more prone to motivational differences. For example, unpublished findings from our lab indicate that birds that pecked for longer in a thwarting task also ate more when the food was available (Verbruggen et al., 2019). Thus, even though all birds were similarly food deprived, it is still possible that some birds were generally more motivated to eat than others, which could in turn cause variation in task performance. However, an advantage of the race model is that, by considering going and stopping together, it may also lend itself to better disentangling the role of cognitive and non-cognitive factors in future research.

## Unravelling Variability Across Species

We observed group differences between lesser black-backed gulls and herring gulls in the detour barrier, thwarting and stop-change tasks. In the detour barrier and stop-change tasks, the lesser black-backed gulls were significantly faster to go and to stop a discrete action than the herring gulls. This faster behaviour was also evident in the stop-change task in how soon they left the start box (relative to the opening of the door), which we used as a general indicator of motivation and activity level. Crucially, the difference in going and stopping persisted even when we considered the species differences in the time taken to leave the start box. In the thwarting task, lesser black-backed gulls were significantly faster to go than the herring gulls, but we did not observe any significant species differences in their stopping behaviour.

The observed behavioural differences in going and stopping might stem from the species’ distinct migration and foraging strategies. For instance, migrant species, such as lesser black-backed gulls, might be less hesitant to explore new environments and be more active in such environments, two traits beneficial during migration (Mettke-Hofmann, 2010, 2014, 2017; Mettke-Hofmann & Gwinner, 2004; Nilsson et al., 2010). This idea aligns with the quicker ‘go’ responses we observed. Furthermore, in the introduction, we speculated that resident species like herring gulls may have adapted to stop discrete actions more efficiently due to the need to adjust their foraging behaviours with seasonal changes. Yet, lesser black-backed gulls were quicker at stopping and changing actions. The speculation that herring gulls might be worse at stopping a repetitive action was also not supported by the data from the detour barrier and thwarting tasks and deserves further research.

Interestingly, in addition to the observed group differences in average go and stopping speed, we found that, across tasks, herring gulls displayed more variation in behaviour between individuals. This might relate to their more ‘generalist’ and variable foraging habits compared to lesser black-backed gulls (Götmark, 1994; McCleery & Sibly, 1986; Pierotti & Annett, 1991; Sotillo et al., 2014; Spaans, 1971; van den Bosch et al., 2019).

In this study, the post-natal environment of the lesser black-backed gulls and herring gulls was standardised. However, we did not have (full) control over their pre-natal environment. Most of our herring gulls came from rooftops, while most of our lesser black-backed gulls came from ground colonies. This may have created different pre-natal environments (e.g. temperature, noise, nutritional values in the yolk, social cues), which could also have influenced cognition and behaviour (Bock et al., 2015).

## Conclusion

In our study, we explored going and stopping behaviour across tasks and species. We found some correlations in the initiation of the going behaviour across tasks, as well as in the stopping behaviour of a discrete action, but less consistency in the stopping behaviour of repetitive actions. The diversity in stopping actions is consistent with previous findings. Despite this, tasks to study the stopping of actions (or even more generally, inhibitory control) are still used interchangeably across studies. This is not only the case in the animal cognition and behavioural ecology domains, but also in the human and (animal) neuroscience domains. Our work and that of others clearly indicates that it is important to consider the various subcomponents of stopping. This is further illustrated by our species comparison: for example, if we had only used a measure of stopping repetitive actions (perseverative pecking at the cover), we would have concluded that lesser black-backed gulls and herring gulls did not significantly differ in stopping (whereas we did observe significant differences in stopping discrete actions). Of course, for practical and ethical reasons, it may not always be possible to include multiple tasks in a study. However, researchers should then select the task most appropriate for their research question and species, provide motivation for this choice (which stopping components are of interest), and, above all, consider the possibility that not all subcomponents of stopping might be equally influenced before generalizing their results. As such, we suggest that researchers are explicit about the type of go and stop stimulus, the timing of the stop stimulus, and type of action to be stopped when using tasks measuring aspects of stopping behaviour, before considering how this might relate to natural behaviour or fitness.

## Acknowledgements

We thank the Gull Patrol (Marc Verborgh, Nathan Noels and Nathalie Colpaert) and ANB for providing the eggs, Michiel Cattrysse (MC) and the VOC staff, particularly Isabelle Allemeersch, Sabine Dedecker, Rijn van Maele, and Claude Velter, for their help with bird care and organisation, Viki Vandomme (VV) and Hans Matheve (HM) for analytical support, Zhang Chen and Maxime Dahirel for help with R coding, and Lies Baten and Angelica Alcantara-Exposito for administrative support.

## Author Contributions (CREDIT)

Conceptualization: C.A.T., A.V., R.A., L.L., F.V.; Methodology: C.A.T., A.V., R.A., L.L., F.V.; Software: F.V.; Investigation: C.A.T., A.V., R.A., S.K., L.L., F.V.; Animal Care: C.A.T., A.V., R.A., L.L., F.V.; Data curation: C.A.T.; Formal analysis: C.A.T.; Visualization: C.A.T.; Validation: A.V. and R.A.; Writing – original draft: C.A.T., L.L., and F.V.; Writing – review and editing: C.A.T, A.V., A.M., L.L., F.V.; Supervision: L.L., and F.V.; Funding acquisition: A.M., L.L., F.V.; Project administration: F.V.

