## Supplementary Material for "Beyond a Unitary Construct: Dissecting Stopping Behaviour in Two Bird Species"

#### 1. Methods

We report our methods following the MeRIT system (Nakagawa et al. 2023).

**Egg collection.** From May 2021 to June 2021, eggs were collected by the Agency for Nature and Forests (ANB) and the Wildlife Rescue Centre (WRC) Ostend, who are authorized to remove gull eggs along the Belgium coast for various reasons. Species was inferred from nest structure or parent species ID. This resulted in a total of 540 herring and lesser black-backed gull eggs from 205 nests along the Belgian coast (De Panne, Oostende, Blankenberge, Zeebrugge, Knokke). These eggs were brought to the WRC on the day of collection.

**Egg incubation.** Upon arrival, CAT, AV and RA weighed, measured and photographed the eggs before placing them in incubators (OVA-Easy Advance; Brinsea; temperature: 37.5°C; humidity until pipping: 45%). For three-egg clutches, only the two largest ones (based on volume) were incubated. Eggs were checked twice a day (around 10:00 and 16:00) for signs of pipping. Pipping eggs were transferred to an incubator with higher humidity for hatching (OVA-Easy Advance; Brinsea; temperature: 37.5°C; humidity 70%). Due to technical issues with the incubators, initial mortality was high and additional eggs were incubated until we reached our target sample size.

**Chick rearing**. CAT, AV, RA, LL, FV and Michiel Cattrysse (MC) reared 120 chicks under standardized conditions. In line with the STRANGE framework, an overview of our sample size is provided in Table S1 (Webster and Rutz 2020). Once hatched, CAT, AV and RA gave chicks a unique combination of coloured rings. For the first 4 days of life, chicks were kept in indoor enclosures (50 x 50 cm), with a heating plate (brand: Comfort). During this time, we hand fed them with smelt, with added vitamins (Akwavit™) and water. Once chicks were 5 days old, and if they weighed more than 60g, they were placed in outside enclosures (1.95 x 5.50 m, covered with pebbles, except for a 1 m long concrete area at the entrance). If they were 5 days old but weighed less than 60g, they were kept in the indoor enclosures until they reached the target weight.

Once outside, chicks were fed mixture of fish (~25%) and puppy food pellets (28.67% protein) soaked in water (~75%) with added vitamins (Akwavit™). Food was delivered in two feeding stations (~1 x 1 m) inside on the concrete area of the enclosure, with a barrier placed at the entrance of the feeding station (Figure 2). Once a group was complete, food was placed in front of the barrier for the first two days. Subsequently and for the remaining of the experiment, food was placed behind the barriers, hence providing individual with the experience of detouring opaque barriers.

**Early-life manipulation.** Four outdoor enclosures of 15 chicks (n=60) were assigned to a 'predictable food' treatment, and four outdoor enclosures of 15 chicks (n=60) were assigned to an 'unpredictable food' treatment. Siblings were split between the two treatments to control for genetic and pre-hatching environment. Species were also split between the two treatments to get an equivalent sample size of each species in each treatment (herring gulls in the predictable treatment: n= 22; herring gulls in the unpredictable treatment: n= 24; lesser black-backed gulls in the predictable treatment: n= 38; lesser black-backed gulls in the unpredictable treatment: n= 36; Table S1). We only knew the sex after the allocation to the treatment, so sex was randomly distributed across treatment (Table S1).

Initially, we aimed to manipulate food predictability for two weeks after group completion. Due to the aforementioned technical issues, we could only implement this scheme for seven days. In brief, time of feeding was either predictable (i.e., feeding was at the same time each day: 9:00, 13:00, 17:00) or unpredictable (i.e., feeding time varied between days: 8:00, 9:00 or 10:00; 12:00, 13:00 or 14:00; 17:00 or 18:00). Furthermore, we added a 'frustration manipulation' (Days 3-7 of the manipulation) during the morning and the evening feeding sessions for chicks in the unpredictable treatment: food that was placed in the enclosure was retrieved for 1 minute as soon as the first chick ate, before giving back the food to the chicks. This procedure was repeated three times during each session, after which the food was left in the enclosure.

**Behavioural tests**. CAT, AV, RA, FV, MC, and LL conducted the behavioural tests. SK coded the videos and RA double coded 20% of the videos. FV extracted distance data in the stop-change task.

**Species ID and sex**. DNA sampling for species ID and sex were conducted by Viki Vandomme, using feather samples collected when chicks hatched. For individuals for which the species could not be identified using DNA methods (e.g., because there were not enough feathers available), RA confirmed the species by using morphological characteristics when individuals were ringed prior to release. The ringing was done by AV and RA. For individuals for which the sex could not be identified using DNA methods, Hans Matheve used a support vector machine classifier to predict individual's sex. Using birds for which we knew the sex and had measurements of weight, tarsus length, wing length, and head length when they were ringed, we predicted the sex of the remaining individuals for each species separately (accuracy of prediction for LBBG: 0.956 [95% C.I. 0.876-0.991], p-value<0.001; n=15; accuracy of prediction for HG: 0.915 [95% C.I. 0.796-0.976], p-value<0.001; n=11). There was one individual for which we did not have molecular data for sex identification, and which died before it was ringed, so we could not sex it using the predictive model. RA and MC measured the tarsus length prior to the behavioural tests.

**Statistical analysis.** CAT conducted the analysis. Using a co-pilot system RA checked the data processing codes, while AV checked the analysis code.

### 2. Data description

Table S1: Following Webster & Rutz (2020), this table provides a description of our sample size based on species, early-life environment, and sex (we were not able to get the sex of one individual).

| | Predictable | Unpredictable |
|---|---|---|
| HG | 22 | 24 |
| LBBG | 38 | 36 |

| | Female | Male |
|---|---|---|
| HG | 27 | 19 |
| LBBG | 34 | 39 |

Table S2: List of individuals that do not do specific behaviour of interest. Highlighted in red are the individuals which were excluded from the analysis (X: did not interact with the task; NA: we do not have the data due to technical issues). 21 individuals did not interact with the detour task, 15 with the thwarting task, and 14 with the stop-change task. In the stop-change task we also lost data for 7 individuals due to technical issues. In total, we had to exclude 33 individuals from our initial analysis.

| | Detour | | | | Thwarting | | | Stop change | | Excluded from correlation and PCA |
|---|---|---|---|---|---|---|---|---|---|---|
| | Did not interact with barrier | Did not detour | Did not eat | Excluded from detour | Did not interact with apparatus | Did not eat | Excluded from thwarting | Did not cross the IR beam (excluded from stop-change) | Did not change | |
| Sample size | 37 | 22 | 28 | 21 | 17 | 31 | 15 | 14 (21 excluded when including NA) | 31 | 33 |
| B_BB | | | | | | | | **X** | X | X |
| B_BR | X | X | X | **X** | | X | | **X** | X | X |
| B_GG | | | | | | X | | | | |
| B_GR | | | | | | X | | | | |
| B_RR | X | | X | **X** | X | X | **X** | | X | X |
| B_RY | X | | | | | | | | | |
| BB_BG | X | X | X | **X** | | X | | **X** | X | X |
| BB_BR | X | X | X | **X** | X | | | **NA** | NA | X |
| BB_R | X | | X | **X** | | | | | | X |
| BBG_GRR | | | | | | | | **NA** | NA | X |
| BG_B | | X | X | | | X | | | | |
| BG_BY | X | | | | | | | | | |
| BG_Y | | | | | | | | | X | |
| BGPR_ | X | | X | **X** | X | X | **X** | | | X |
| BR_BB | X | | | | | | | | | |

| | | | | | | | | | | |
|---|---|---|---|---|---|---|---|---|---|---|
| BR_RY | X | X | X | **X** | X | X | **X** | **X** | X | X |
| BR_Y | | | | | | X | | | | |
| BY_RY | | | | | | X | | | | |
| BY_YY | X | X | X | **X** | X | X | **X** | **X** | X | X |
| BYY_ | | | | | | | | | X | |
| G_BB | X | | | | | | | | | |
| G_BG | | X | X | | | X | | | X | |
| G_GR | | | | | | | | | X | |
| G_GY | | | | | X | X | **X** | | X | X |
| G_RR | | | | | | X | | | | |
| G_RY | | | | | | | | | X | |
| GG_BB | | | | | | | | **NA** | NA | X |
| GG_BP | X | X | X | **X** | X | X | **X** | **X** | X | X |
| GG_BR | X | | | | | | | | | |
| GG_GG | X | | | | | | | | | |
| GP_B | X | | | | | | | | | |
| GP_BB | X | X | X | **X** | X | X | **X** | | | X |
| GP_BG | X | | | | | | | | | |
| GP_GY | | X | X | | | | | | | |
| GP_R | X | | | | | | | | | |
| GP_YY | X | X | X | **X** | | X | | | | X |
| GPY_Y | X | X | X | **X** | X | | | | X | X |
| GY_B | | | | | | X | | | X | |
| GY_BR | X | | | | | | | | | |
| P_B | | | | | | | | **NA** | NA | X |
| P_BB | X | X | X | **X** | X | X | **X** | **X** | X | X |
| P_BY | | | | | | X | | | | |
| P_GY | | | | | | X | | | X | |
| P_P | | | | | | X | | | | |

| | | | | | | | | | | |
|---|---|---|---|---|---|---|---|---|---|---|
| P_Y | | | | | | | | **X** | X | X |
| PR_PP | X | | | | | | | **NA** | NA | X |
| PR_Y | X | | | | X | X | **X** | | | X |
| PY_YY | | | | | | | | | X | |
| R_BG | X | | | | | | | | | |
| R_BGY | X | | | | | | | | | |
| R_BPR | | | | | | X | | | X | |
| R_BR | | X | X | | | | | | X | |
| R_BRR | X | X | X | **X** | X | X | **X** | | | X |
| R_GG | X | X | X | **X** | | | | | | X |
| R_GR | X | X | X | **X** | X | X | **X** | **X** | X | X |
| R_GRR | X | | X | **X** | X | X | **X** | **X** | X | X |
| R_GY | | | | | | | | **NA** | NA | X |
| R_R | | X | X | | | | | | X | |
| RR_BY | | | | | | X | | | | |
| RY_BY | X | X | X | **X** | X | X | **X** | **X** | X | X |
| RY_G | | X | X | | | | | | X | |
| RY_P | | | X | | X | X | **X** | | X | X |
| Y_B | X | | | | | | | | | |
| Y_BY | | | | | | | | **X** | X | X |
| Y_G | X | X | X | **X** | | | | | | X |
| Y_GR | X | X | X | **X** | | | | **X** | X | X |
| Y_P | | | | | | | | **NA** | NA | X |
| YY_BB | X | | | | | | | | X | |
| YY_P | X | | X | **X** | X | X | **X** | **X** | X | X |

#### 3. Factors influencing participation (exclusion criteria)

We first examined whether participation (whether the bird participated (1) or not (0)) was repeatable across all three tasks using the rptR package (version 0.9.22, Stoffel, Nakagawa, and Schielzeth 2017) and a Binary distribution, and found that it was (r = 8.44, CI = 16.8,101.5, p<0.001).

In order to verify whether there was any bias in participation in the tasks, we used a binomial model distribution from the *lmerTest* package (version 3.1.3, Kuznetsova, Brockhoff, and Christensen 2017). We used a separate model for each of the three tasks. Whether a bird participated (1) or not (0) was our dependent variable (model family = binomial). We used species, treatment, sex and tarsus as fixed effects and enclosure as a random effect. One individual was excluded because we did not know its sex (n=119). We did not find any significant effects of either of our fixed effects on participation in each of the three tasks (Tables S3-5).

##### a. Detour barrier task

Table S3: Model showing the effect of species, treatment, sex, and average tarsus length on whether a bird participated or not in the detour barrier task (n=119). Enclosure was included as a random effect. [1] baseline = herring gull; [2] baseline = predictable treatment; [3] baseline = female

| Variable | Estimate (std. error) | 95% CI | P value |
|---|---|---|---|
| **Intercept** | **1.46 (0.523)** | **0.487; 2.59** | **0.005** |
| Species[1]: LBBG | -0.352 (0.534) | -1.45; 0.670 | 0.510 |
| Treatment[2]: unpredictable | 0.124 (0.520) | -0.948; 1.34 | 0.812 |
| Sex[3]: male | 0.965 (0.592) | -0.149; 2.21 | 0.103 |
| Tarsus length | 0.424 (0.265) | -0.097; 0.973 | 0.111 |

##### b. Thwarting task

Table S4: Model showing the effect of species, treatment, sex, and average tarsus length on whether a bird participated or not in the thwarting task (n=119). Enclosure was included as a random effect. [1] baseline = herring gull; [2] baseline = predictable treatment; [3] baseline = female

| Variable | Estimate (std. error) | 95% CI | P value |
|---|---|---|---|
| **Intercept** | **1.67 (0.828)** | **-0.023; 3.99** | **0.044** |
| Species[1]: LBBG | 0.626 (0.623) | -0.599; 1.89 | 0.315 |
| Treatment[2]: unpredictable | 0.717 (1.05) | -1.73; 3.76 | 0.496 |
| Sex[3]: male | 0.300 (0.692) | -1.05; 1.72 | 0.665 |
| Tarsus length | 0.549 (0.346) | -0.120; 1.29 | 0.113 |

##### c. Stop-change task

Table S5: Model showing the effect of species, treatment, sex, and average tarsus length on whether a bird participated or not in the stop-change task (n=112). Enclosure was included as a random effect. [1] baseline = herring gull; [2] baseline = predictable treatment; [3] baseline = female

| Variable | Estimate (std. error) | 95% CI | P value |
|---|---|---|---|

| | | | |
|---|---|---|---|
| **Intercept** | **2.05 (0.630)** | **0.929; 3.52** | **0.001** |
| Species[1]: LBBG | -0.366 (0.645) | -1.74; 0.849 | 0.571 |
| Treatment[2]: unpredictable | 0.491 (0.613) | -0.797; 1.85 | 0.423 |
| Sex[3]: male | 0.028 (0.673) | -1.30; 1.38 | 0.967 |
| Tarsus length | 0.134 (0.325) | -0.558; 0762 | 0.679 |

### 4. Inter-coder reliability

Table S6: Table showing for each behavioural variable used in the analysis, the interclass correlation coefficient (ICC), its p-value, and the sample size of the comparison between the two ratters (i.e. number of videos that were double-coded in which the behaviour occurred)

| Task | Variable Inclusion | Measure | Sample size | ICC | p-value |
|---|---|---|---|---|---|
| Detour | Analysis | Time spent interacting with the barrier | 21 | 0.992 | <0.001 |
| | | Latency to exit | 24 | 0.999 | <0.001 |
| | | Latency to detour | 21 | 1.00 | <0.001 |
| | Analysis and Data inclusion | Latency to interact with the barrier | 21 | 0.907 | <0.001 |
| | Data inclusion | Latency to eat | 20 | 1 | <0.001 |
| Thwarting | Analysis | Time spent interacting with the apparatus | 20 | 0.971 | <0.001 |
| | | Latency to exit | 24 | 1.00 | <0.001 |
| | Analysis and Data inclusion | Latency to interact with the apparatus | 20 | 0.962 | <0.001 |
| | Data inclusion | Latency to eat | 17 | 0.998 | <0.001 |
| Stop-change | Analysis | Latency to exit | 23 | 0.997 | <0.001 |
| | | Latency to change | 15 | 0.993 | <0.001 |
| | Analysis and Data inclusion | Latency to cross the IR beam | 18 | 0.989 | <0.001 |

### 5. Analyses only including birds that interact with the barrier in the detour task

For the detour task, we defined the latency to interact with the apparatus as the latency to interact with the barrier, or for those that did not interact with the barrier (n=16) the latency to detour. The correlation between the latency to interact with the apparatus and the latency to detour may have been spuriously increased as a result (as for 16 birds this was the same measure). We therefore reran the correlation between all variables (Table S7, Figure S1; n=73) and the PCA (Table S8; n=73) and the GLMM looking at the latency to interact with the detour barrier task (Table S9-10; n=83), by excluding those birds. The results are reported below and the main results remain unchanged.

#### a. Correlations and PCA (n=73)

Table S7: Correlation matrix showing the correlation coefficient and Bayes factor between the different behavioural measures (N=87). In bold are results supporting moderate to extreme evidence for the alternative hypothesis, and in italic are results supporting moderate evidence for the null hypothesis. In (A) are behavioural measures corresponding to the “go” component; in (B) are those corresponding to the “stopping” component.

| A. Going | Detour barrier: latency to interact with the task | Thwarting: Latency to interact with task |
|---|---|---|
| Thwarting: latency to interact with task | 0.257 (2.52) | |
| Stop-change: latency to interact task | 0.246 (2.08) | 0.090 (0.349) |

| B. Stopping | Detour barrier: Latency to detour | Detour barrier: time spent interacting with the barrier | Thwarting: time spent interacting with the apparatus | Stop-change: latency to change |
|---|---|---|---|---|
| Detour barrier: time spent interacting with the barrier | **0.523 (9300)** | | | |
| Thwarting: time spent interacting with the apparatus | -0.186 (0.846) | *0.075 (0.322)* | | |
| Stop-change: latency to change | **0.278 (3.75)** | -0.125 (0.446) | -0.216 (1.28) | |
| Stop-change: minimum distance to old location | 0.124 (0.443) | *-0.033 (0.277)* | -0.259 (2.61) | **0.374 (37.5)** |

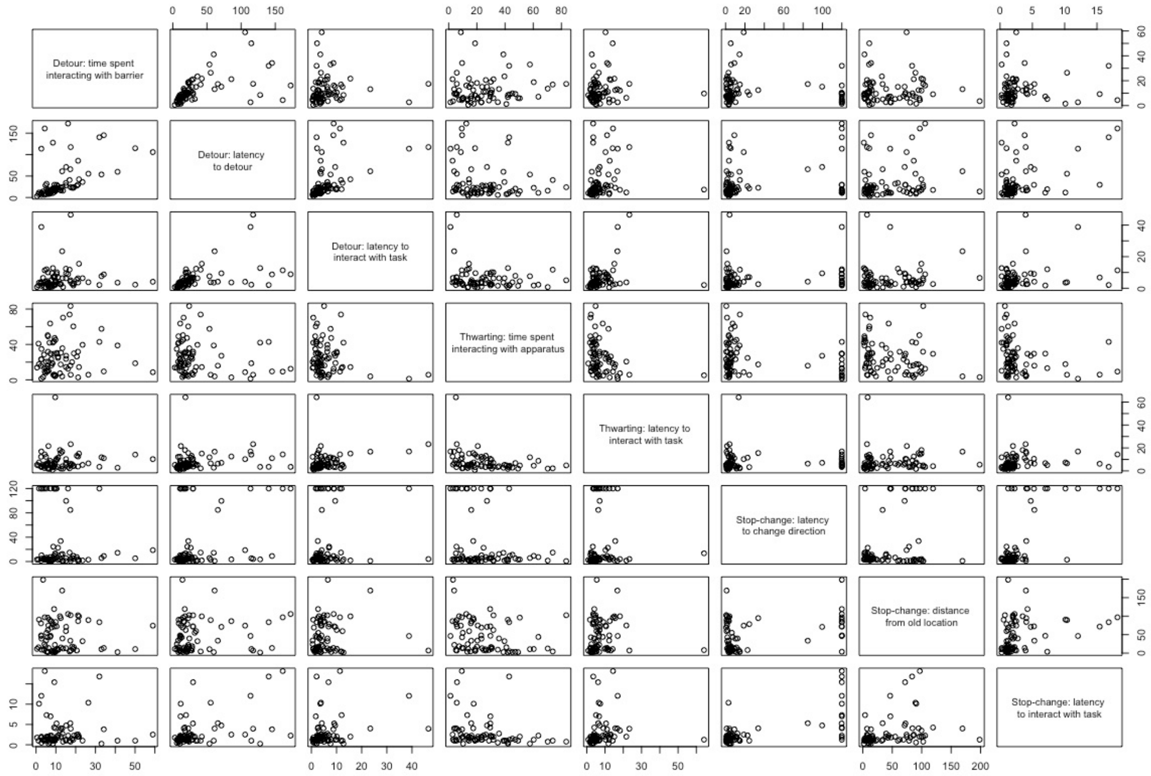


<u>Figure S1:</u> Correlation plot between the different behavioural measures used in all three tasks.

<u>Table S8:</u> Structure of the principal component analysis for behavioural measures related to stopping behaviour in the detour, thwarting and stop-change tasks. In bold we report the loading for variables whose loading is bigger than expected if all variables contributed equally to the specific axis.

| | Axis 1 | Axis 2 | Axis 3 | Axis 4 | Axis 5 | Axis 6 | Axis 7 | Axis 8 |
|---|---|---|---|---|---|---|---|---|
| Std deviation | 1.60 | 1.23 | 1.10 | 0.930 | 0.834 | 0.784 | 0.555 | 0.494 |
| Proportion of variance | 0.320 | 0.188 | 0.151 | 0.108 | 0.087 | 0.077 | 0.039 | 0.030 |
| Cumulative proportion | 0.320 | 0.508 | 0.659 | 0.767 | 0.854 | 0.931 | 0.970 | 1.00 |
| Detour: latency to interact with the task (Go) | -0.347 | 0.238 | 0.272 | **-0.567** | **0.529** | -0.141 | -0.061 | -0.352 |
| Thwarting: latency to interact with the apparatus (Go) | -0.212 | 0.204 | **0.634** | 0.165 | **-0.462** | **-0.513** | -0.010 | 0.087 |
| Stop-change: latency to interact with the task (Go) | **-0.489** | -0.186 | -0.244 | -0.167 | -0.342 | 0.032 | **0.691** | -0.209 |
| Detour: latency to detour (Stop) | **-0.437** | **0.450** | -0.224 | -0.058 | 0.077 | 0.119 | -0.029 | **0.729** |
| Detour: time spent interacting with the barrier (Stop) | -0.108 | **0.637** | -0.320 | 0.431 | -0.085 | 0.031 | -0.116 | **-0.522** |

| | | | | | | | | |
|---|---|---|---|---|---|---|---|---|
| Thwarting: time spent interacting with the apparatus (Stop) | 0.331 | 0.119 | **-0.486** | -0.299 | -0.043 | **-0.735** | 0.059 | 0.078 |
| Stop-change: latency to change (Stop) | **-0.433** | -0.371 | -0.264 | -0.124 | -0.287 | -0.081 | **-0.700** | -0.106 |
| Stop-change: minimum distance to old location (Stop) | -0.308 | **-0.327** | -0.076 | **0.575** | **0.541** | **-0.393** | 0.107 | 0.051 |

#### b. Detour: latency to interact with the apparatus (n=83)

Table S9: Model showing the effect of species, treatment, sex, latency to exit, and tarsus length on the latency to interact with the apparatus in the detour task (n=83). This only includes bird that did interact with the barrier, but not those that detoured without interacting with the barrier (n=16). Enclosure was included as a random effect. Based on the model residual diagnostics, the negative binomial model accounting for overdispersion was the best fit. The dispersion model included species, treatment and sex as fixed effects. [1] baseline = herring gull; [2] baseline = predictable treatment; [3] baseline = female

| Variable | Estimate (std. error) | 95% CI | P value |
|---|---|---|---|
| **Intercept** | **2.11 (0.189)** | **1.74; 2.49** | **<0.001** |
| **Species[1]: LBBG** | **-0.706 (0.166)** | **-1.03; -0.380** | **<0.001** |
| Treatment[2]: unpredictable | -0.273 (0.171) | -0.608; 0.061 | 0.109 |
| Sex[3]: male | 0.257 (0.170) | -0.076; 0.589 | 0.131 |
| **Latency to exit** | **0.171 (0.080)** | **0.014; 0.327** | **0.032** |
| Tarsus length | 0.016 (0.081) | -0.143; 0.175 | 0.843 |

Table S10: Dispersion model showing the effect of species, treatment and sex on the dispersion parameter, for the model looking at the latency to interact with the apparatus in the detour task. [1] baseline = herring gull; [2] baseline = predictable treatment; [3] baseline = female

| Variable | Estimate (std. error) | 95% CI | P value |
|---|---|---|---|
| **Intercept** | **1.81 (0.456)** | **0.915; 2.70** | **<0.001** |
| **Species[1]: LBBG** | **-1.83 (0.670)** | **-3.14; -0.512** | **0.006** |
| **Treatment[2]: unpredictable** | **-1.68 (0.650)** | **-2.95; -0.402** | **0.010** |
| Sex[3]: male | 0.893 (0.592) | -0.268; 2.05 | 0.132 |

### 6. Stopping behaviour across tasks (n=87)

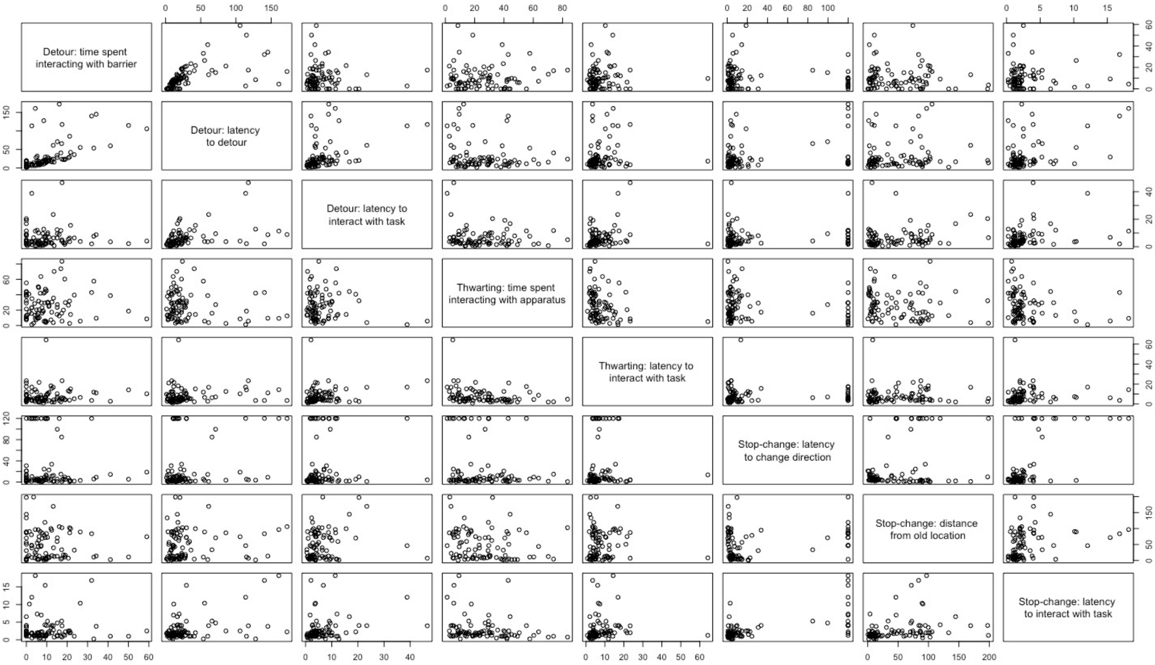


Figure S2: Correlation plot between the different behavioural measures used in all three tasks.

Table S11: Structure of the principal component analysis for behavioural measures related to stopping behaviour in the detour, thwarting and stop-change task. In bold we report the loading for variables whose loading is bigger than expected if all variables contributed equally to the specific axis.

| | Axis 1 | Axis 2 | Axis 3 | Axis 4 | Axis 5 | Axis 6 | Axis 7 | Axis 8 |
|---|---|---|---|---|---|---|---|---|
| Std deviation | 1.56 | 1.21 | 1.09 | 0.941 | 0.900 | 0.812 | 0.579 | 0.472 |
| Proportion of variance | 0.303 | 0.184 | 0.149 | 0.111 | 0.101 | 0.082 | 0.042 | 0.027 |
| Cumulative proportion | 0.303 | 0.487 | 0.636 | 0.747 | 0.848 | 0.930 | 0.972 | 1.00 |
| Detour: latency to interact with the task (Go) | 0.351 | -0.044 | -0.270 | **0.781** | -0.060 | 0.243 | -0.132 | 0.335 |
| Thwarting: latency to interact with the apparatus (Go) | 0.181 | -0.085 | **-0.699** | -0.135 | -0.327 | **-0.583** | 0.036 | -0.069 |
| Stop-change: latency to interact with the task (Go) | **0.511** | 0.164 | 0.249 | -0.118 | -0.253 | 0.028 | **0.732** | 0.186 |
| Detour: latency to detour (Stop) | **0.462** | **-0.477** | 0.101 | -0.067 | 0.058 | 0.129 | -0.041 | **-0.723** |
| Detour: time spent interacting with the task (Stop) | 0.146 | **-0.702** | 0.161 | -0.210 | 0.259 | -0.213 | -0.074 | **0.546** |
| Thwarting: time spent interacting with the apparatus (Stop) | -0.275 | -0.139 | **0.487** | **0.479** | **-0.356** | **-0.545** | 0.058 | -0.100 |

| | | | | | | | | |
|---|---|---|---|---|---|---|---|---|
| Stop-change: latency to change (Stop) | **0.438** | 0.283 | 0.311 | -0.233 | -0.351 | -0.082 | **-0.659** | 0.110 |
| Stop-change: minimum distance to old location (Stop) | 0.283 | **0.380** | 0.075 | 0.158 | **0.711** | **-0.484** | 0.001 | -0.074 |

7. **Factors influencing stopping behaviour**
    a. **Selection in model families**

Because all of our behaviours of interest were count data (time or distance), we started with using the Poisson family model. Based on the diagnostics from *DHARMa*, if model assumptions were violated, we then tried negative binomial distribution. If the model diagnostics showed evidence of zero-inflation or heteroskedasticity, we also included a zero-inflated model and a dispersion model. For the latency to interact with the detour task, the final model family was a negative binomial model accounting for overdispersion (Table 4.A). The dispersion model included species, treatment, and sex (Table S12). For the latency to interact with the thwarting task, the final model family was a negative binomial model accounting for overdispersion (Table 4.B). The dispersion model included species, treatment, and sex (Table S15). For the latency to interact with the stop-change task, the final model family was a negative binomial model accounting for overdispersion (Table 4.C). The dispersion model included species, and sex (Table S16). For the latency to detour in the detour barrier task, the final model family was a negative binomial model accounting for overdispersion (Table 4.D). The dispersion model included species, treatment, and sex (Table S13). For the latency to change direction in the stop-change task, the final model family was a negative binomial model accounting for overdispersion (Table 4.E). The fixed effects of the dispersion model included species, treatment, and sex (Table S14). For the time spent interacting with the barrier in the detour barrier task, the final model family was a zero-inflated negative binomial model (Table S17). The fixed effects of the zero-inflated model were the same as in the main model (Table S18). For the time spent interacting with the apparatus in the thwarting task, the final model family was a negative binomial model (Table S19). For the minimum distance from old location in the stop-change task, the final model family was a negative binomial model (Table S20).

    b. **Detour: latency to interact with the apparatus (n=99)**

Table S12: Dispersion model showing the effect of species, treatment and sex on the dispersion parameter, for the model looking at the latency to interact with the apparatus in the detour barrier task. [1] baseline = herring gull; [2] baseline = predictable treatment; [3] baseline = female

| Variable | Estimate (std. error) | 95% CI | P value |
|---|---|---|---|
| **Intercept** | **1.90 (0.422)** | **1.08; 2.73** | **<0.001** |
| **Species[1]: LBBG** | **-1.24 (0.532)** | **-2.29; -0.197** | **0.020** |
| Treatment[2]: unpredictable | -0.694 (0.480) | -1.07; 0.715 | 0.149 |
| Sex[3]: male | -0.180 (0.456) | 0.009; 1.30 | 0.693 |

c. **Detour: latency to detour (n=93)**

Table S13: Dispersion model showing the effect of species, treatment and sex on the dispersion parameter, for the model looking at the latency to detour in the detour barrier task. [1] baseline = herring gull; [2] baseline = predictable treatment; [3] baseline = female

| Variable | Estimate (std. error) | 95% CI | P value |
|---|---|---|---|
| **Intercept** | **3.58 (0.341)** | **2.91; 4.25** | **<0.001** |
| **Species[1]: LBBG** | **-1.19 (0.378)** | **-1.93; -0.445** | **0.002** |
| **Treatment[2]: unpredictable** | **-1.37 (0.391)** | **-2.14; -0.601** | **<0.001** |
| Sex[3]: male | -0.974 (0.373) | -0.613; 0.849 | 0.752 |

d. **Stop-change: latency to change direction (n=82)**

Table S14: Dispersion model showing the effect of species, treatment and sex on the dispersion parameter, for the model looking at the latency to change direction in the stop-change task. [1] baseline = herring gull; [2] baseline = predictable treatment; [3] baseline = female

| Variable | Estimate (std. error) | 95% CI | P value |
|---|---|---|---|
| **Intercept** | **3.04 (0.533)** | **2.00; 4.08** | **<0.001** |
| **Species[1]: LBBG** | **-2.12 (0.457)** | **-3.02; -1.23** | **<0.001** |
| Treatment[2]: unpredictable | 0.325 (0.430) | -0.518; 1.17 | 0.450 |
| Sex[3]: male | 0.105 (0.460) | -0.796; 1.01 | 0.819 |

e. **Thwarting: latency to interact with the apparatus (n=103)**

Table S15: Dispersion model showing the effect of species, treatment and sex on the dispersion parameter, for the model looking at the latency to interact with the apparatus in the thwarting task. [1] baseline = herring gull; [2] baseline = predictable treatment; [3] baseline = female

| Variable | Estimate (std. error) | 95% CI | P value |
|---|---|---|---|
| **Intercept** | **1.21 (0.450)** | **0.332; 2.09** | **0.007** |
| Species[1]: LBBG | 0.215 (0.432) | -0.632; 1.06 | 0.619 |
| Treatment[2]: unpredictable | 0.291 (0.429) | -0.550; 1.13 | 0.498 |
| **Sex[3]: male** | **-1.32 (0.454)** | **-2.21; -0.427** | **0.004** |

f. **Stop-change: latency to interact with the apparatus (trigger the seesaw) (n=98)**

Table S16: Dispersion model showing the effect of species, treatment and sex on the dispersion parameter, for the model looking at the latency to interact with the stop-change task. [1] baseline = herring gull; [2] baseline = female

| Variable | Estimate (std. error) | 95% CI | P value |
|---|---|---|---|
| Intercept | 0.651 (0.587) | -0.500; 1.80 | 0.268 |
| Species[1]: LBBG | -1.88 (1.42) | -4.67; 0.917 | 0.188 |
| Treatment[2]: unpredictable | -0.200 (1.08) | -2.32; 1.92 | 0.853 |
| Sex[2]: male | -3.264 (7.09) | -17.2; 10.6 | 0.645 |

#### g. Detour: time spent interacting with the detour barrier (n=99)

Table S17: Model showing the effect of species, treatment, sex and latency to exit on the time spent interacting with the barrier in the detour barrier task (n=99). Enclosure was included as a random effect. Based on the model residual diagnostics, the zero-inflated negative binomial model was the best fit. The zero-inflated model had the same fixed effects as the main model. [1] baseline = herring gull; [2] baseline = predictable treatment; [3] baseline = female

| Variable | Estimate (std. error) | 95% CI | P value |
|---|---|---|---|
| **Intercept** | **2.90 (0.156)** | **2.60; 3.21** | **<0.001** |
| Species[1]: LBBG | -0.234 (0.165) | -0.558; 0.091 | 0.158 |
| Treatment[2]: unpredictable | -0.257 (0.176) | -0.602; 0.088 | 0.144 |
| Sex[3]: male | -0.239 (0.156) | -0.544; 0.066 | 0.125 |
| Latency to exit | -0.028 (0.092) | -0.207; 0.151 | 0.759 |

Table S18: Zero-inflated model showing the effect of species, treatment and sex on the zero-inflation parameter, for the model looking at the time spent interacting with the barrier in the detour barrier task. [1] baseline = herring gull; [2] baseline = predictable treatment; [3] baseline = female

| Variable | Estimate (std. error) | 95% CI | P value |
|---|---|---|---|
| **Intercept** | **-1.24 (0.582)** | **-2.38; -0.095** | **0.034** |
| Species[1]: LBBG | 0.612 (0.675) | -0.711; 1.94 | 0.365 |
| Treatment[2]: unpredictable | -1.12 (0.711) | -2.52; 0.270 | 0.114 |
| Sex[3]: male | -0.974 (0.689) | -2.32; 0.376 | 0.157 |
| Latency to exit | 0.515 (0.272) | -0.019; 1.05 | 0.059 |

#### h. Thwarting: time spent interacting with the apparatus (n=105)

Table S19: Model showing the effect of species, treatment, sex and latency to exit on the time spent interacting with the apparatus in the thwarting task (n=105). Enclosure was included as a random effect. Based on the model residual diagnostics, the negative binomial model was the best fit. [1] baseline = herring gull; [2] baseline = predictable treatment; [3] baseline = female

| Variable | Estimate (std. error) | 95% CI | P value |
|---|---|---|---|
| **Intercept** | **3.11 (0.157)** | **2.81; 3.43** | **<0.001** |
| Species[1]: LBBG | -0.195 (0.122) | -0.434; 0.044 | 0.110 |
| Treatment[2]: unpredictable | 0.287 (0.174) | -0.055; 0.628 | 0.100 |
| Sex[3]: male | -0.011 (0.117) | -0.241; 0.219 | 0.923 |
| **Latency to exit** | **-0.498 (0.088)** | **-0.671; -0.324** | **<0.001** |

#### i. Stop-change: minimum distance to the old location (n=99)

Table S20: Model showing the effect of species, treatment, sex, latency to exit and average tarsus length on the minimum distance to the old location (n=99). Enclosure was included as a random effect. Based on the model residual diagnostics, the negative binomial model was the best fit. [1] baseline = herring gull; [2] baseline = predictable treatment; [3] baseline = female

| Variable | Estimate (std. error) | 95% CI | P value |
|---|---|---|---|
| **Intercept** | **3.61 (0.178)** | **3.27; 3.96** | **<0.001** |
| Species[1]: LBBG | 0.031 (0.163) | -0.288; 0.351 | 0.848 |
| **Treatment[2]: unpredictable** | **0.442 (0.165)** | **0.118; 0.765** | **0.007** |
| Sex[3]: male | -0.020 (0.176) | -0.364; 0.325 | 0.911 |
| **Latency to exit** | **0.212 (0.071)** | **0.073; 0.351** | **0.003** |
| Tarsus length | -0.029 (0.097) | -0.219; 0.160 | 0.763 |

### 8. Factors influencing latency to exit the start box

Given that there were some evidence of species differences on latency measures in our task, we wanted to check whether those differences could be explained by a more general difference in speed between those two species. We first examined whether latencies to exit the start box were repeatable across all three tasks using the rptR package (version 0.9.22, Stoffel, Nakagawa, and Schielzeth 2017) and a Poisson distribution (after rounding the latencies).

To estimate the effect of species on latency to exit, we used a negative binomial model and zero-inflated models from the *lmerTest* package (version 3.1.3, Kuznetsova, Brockhoff, and Christensen 2017) with latency to exit the start box in each of the three tasks as dependent variables, species, treatment, sex and tarsus as fixed effects and enclosure as a random effect. We used a separate model for each of the three tasks, and only included birds that participate in the task. In the detour and thwarting task, we found weak but non-significant evidence that lesser black backed gulls were faster at exiting the start box than herring gulls (Tables S21-S24), while in the stop-change task we found significant evidence that lesser black backed gulls were faster at exiting the start box than herring gulls (Tables S25).

#### a. Detour barrier task

Table S21: Model showing the effect of species, treatment, sex, and average tarsus length on the latency to exit in the detour task (n=99). Enclosure was included as a random effect. Based on the model residual diagnostics, the negative binomial model accounting for overdispersion was the best fit. [1] baseline = herring gull; [2] baseline = predictable treatment; [3] baseline = female

| Variable | Estimate (std. error) | 95% CI | P value |
|---|---|---|---|
| **Intercept** | **3.00 (0.239)** | **2.53; 3.47** | **<0.001** |
| Species[1]: LBBG | -0.474 (0.261) | -0.986; 0.037 | 0.069 |
| Treatment[2]: unpredictable | 0.079 (0.278) | -0.466; 0.623 | 0.777 |
| Sex[3]: male | 0.037 (0.272) | -0.497; 0.570 | 0.892 |
| Tarsus length | -0.112 (0.096) | -0.300; 0.075 | 0.241 |

Table S22: Dispersion model showing the effect of species, treatment and sex on the dispersion parameter, for the model looking at the latency to exit the start box in the detour task. [1] baseline = herring gull; [2] baseline = predictable treatment; [3] baseline = female

| Variable | Estimate (std. error) | 95% CI | P value |
|---|---|---|---|
| **Intercept** | **3.06 (0.387)** | **2.30; 3.82** | **<0.001** |

| | | | |
|---|---|---|---|
| Species[1]: LBBG | -0.348 (0.415) | -1.16; 0.466 | 0.403 |
| Treatment[2]: unpredictable | 0.242 (0.450) | -0.640; 1.12 | 0.591 |
| Sex[3]: male | -0.012 (0.427) | -0.848; 0.824 | 0.977 |

#### b. Thwarting task

Table S23: Model showing the effect of species, treatment, sex, and average tarsus length on the latency to exit in the thwarting task (n=105). Enclosure was included as a random effect. Based on the model residual diagnostics, the negative binomial model accounting for overdispersion was the best fit. [1] baseline = herring gull; [2] baseline = predictable treatment; [3] baseline = female

| Variable | Estimate (std. error) | 95% CI | P value |
|---|---|---|---|
| **Intercept** | **2.95 (0.242)** | **2.48; 3.43** | **<0.001** |
| Species[1]: LBBG | -0.476 (0.243) | -0.952; 0.001 | 0.051 |
| Treatment[2]: unpredictable | 0.272 (0.244) | -0.207; 0.751 | 0.266 |
| Sex[3]: male | -0.170 (0.253) | -0.666; 0.327 | 0.503 |
| Tarsus length | 0.100 (0.097) | -0.090; 0.290 | 0.304 |

Table S24: Dispersion model showing the effect of species, treatment and sex on the dispersion parameter, for the model looking at the latency to exit the start box in the thwarting task. [1] baseline = herring gull; [2] baseline = predictable treatment; [3] baseline = female

| Variable | Estimate (std. error) | 95% CI | P value |
|---|---|---|---|
| **Intercept** | **3.00 (0.373)** | **2.27; 3.73** | **<0.001** |
| Species[1]: LBBG | -0.196 (0.372) | -0.924; 0.533 | 0.599 |
| Treatment[2]: unpredictable | 0.415 (0.372) | -0.313; 1.14 | 0.264 |
| Sex[3]: male | -0.185 (0.356) | -0.882; 0.513 | 0.603 |

#### c. Stop-change task

Table S25: Model showing the effect of species, treatment, sex, and average tarsus length on the latency to exit in the stop-change task (n=99). Enclosure was included as a random effect. Based on the model residual diagnostics, the = negative binomial model was the best fit. [1] baseline = herring gull; [2] baseline = predictable treatment; [3] baseline = female

| Variable | Estimate (std. error) | 95% CI | P value |
|---|---|---|---|
| **Intercept** | **2.30 (0.206)** | **1.90; 2.70** | **<0.001** |
| **Species[1]: LBBG** | **-0.430 (0.208)** | **-0.837; -0.023** | **0.038** |
| Treatment[2]: unpredictable | 0.194 (0.192) | -0.183; 0.570 | 0.313 |
| Sex[3]: male | 0.236 (0.224) | -0.223; 0.695 | 0.314 |
| Tarsus length | -0.010 (0.111) | -0.228; 0.208 | 0.928 |
